# A broad-spectrum, biocompatible, virucidal polymer reduces chikungunya virus in murine models

**DOI:** 10.1101/2025.10.27.684922

**Authors:** Shannan-Leigh Macleod, Anna X.Y. Loo, Maria A. Castillo, Lauren J. Batt, Fok-Moon Lum, Anthony Torres-Ruesta, Lisa F.P. Ng, Samuel T. Jones

## Abstract

Autochthonous transmission of arboviruses poses significant threats to global health and economies. Yet, no effective antivirals exist. Building on our previous antiviral star-polymer, we designed zwitterionic star-polymers for efficacy in high protein environments. A polymer with 12% positively charged monomer (Zwitterionic Polymer-ZP12) exhibited broad-spectrum, biocompatible antiviral activity against *Alphaviridae*, *Flaviviridae*, *Herpesviridae*, and *Picornaviridae*. Using murine models for Chikungunya virus (CHIKV) infection, ZP12 treatment (10 mg/kg every 24 hours for 7 days) reduced tissue viral load by 90% 3 days post-infection and significantly alleviated CHIKV-induced joint swelling. Mechanistically, ZP12 downregulated CHIKV-driven immunopathogenesis by reducing viral load and dampening CD4**^+^** T cell and macrophage activation in virus-infected joints. With no current antiviral interventions for these arboviruses, ZP12 represents a promising intervention for combating future pandemics.

## 1 Introduction

Arboviruses, viruses that are transmitted by arthropods, such as mosquitoes, are predicted to be the causative agent of the next global pandemic[1]. Global infection rates of arboviruses, such as dengue viruses, yellow fever virus and Zika virus, are rising annually, further increasing the likelihood of future pandemics. Autochthonous transmission of arboviruses to non-tropical regions has also been observed recently due to the expansion of the natural range of virus-associated vectors (mosquitoes[2–6], ticks[7–9] etc.) directly linked with changes in climate, globalisation and urbanisation[10–13]. Chikungunya virus (CHIKV), a tropical arbovirus transmitted to humans by *Aedes* mosquitoes[3–5], is the aetiological agent for Chikungunya fever (CHIKF). CHIKF is characterised by prolonged incapacitating polyarthralgia, with patients experiencing joint pain and inflammation; a high fever (*>*38^◦^C) and myalgia (muscle pain)[10]. Despite infections going largely unreported, approximately 460,000 cases were recorded in 2024 in over 118 countries worldwide[14, 15]. Currently, there are no approved antiviral therapeutics effective against CHIKV or other arboviruses[2, 16, 17].

Small molecule drugs[18–23], natural extracts[17, 24], and oligomers[25, 26], targeting the CHIKV replication cycle, have been extensively studied. Even though CHIKV infection has been reduced in both *in vitro* and *in vivo* models, these strategies rely on specific sequences (*i.e.* DNA, amino acids, antigens etc.) and upon viral mutational drifts, many were deemed ineffective[21, 27, 28]. For wide-ranging broad-spectrum protection against CHIKV (and other viruses) alternative antiviral mechanisms, in particular extracellular modes of action, need to be investigated[9, 16, 29–32].

Like many viruses, CHIKV predominantly uses cell-surface heparan sulfate proteoglycans (HSPGs) as attachment factors for entry into host cells[9, 33–36]. Developing HSPG mimics has been explored as a route to prevent the attachment step of viral replication and may consequently pose broad-spectrum properties[30, 32, 37, 38]. Sulfonated polymers, including heparin[29, 39], are effective HSPG mimics which have previously demonstrated broad-spectrum antiviral properties against several different viruses[9, 37, 40]. However, upon local changes in environment (*i.e.* dilution, pH, conductivity), such polymers typically dissociate from the virus, releasing infectious virions. This renders them virustatic and therefore limits their clinical efficacy[38, 41–43]. Ideally, a broad-spectrum and irreversible virucidal inhibitory mechanism, that alters the integrity of the virions, needs to be developed to prevent the loss of efficacy upon dilution[29, 39, 44, 45]. However, previously identified virucidal materials, such as bleach are toxic, which means that they cannot be used as a treatment option[30].

Recent studies show that increased polyvalency of sulfonated polymers improved virucidal properties, demonstrated with nanoparticles[31, 44, 46–49], cyclodextrins[29, 31, 50–52], and non-linear polymers[40, 53–55]. We have previously shown that a star-polymer (referred to as star-PSS)[45] was a highly efficacious virucidal antiviral when used for *in vivo* topical treatment of viral infections, specifically via intranasal and intravaginal delivery, where the site of action, *i.e.* the lungs and the vagina, had a low protein environment[45]. However, in high protein biological fluids (such as blood, particularly important for arboviruses), highly negatively charged sulfonated moieties have been shown to be coated by a protein corona, significantly reducing their antiviral efficacy[37, 56–60]. This is potentially a problem for systemic viruses, such as CHIKV, as the extracellular antiviral needs to be active in a high protein environment[61–64]. If this promising class of broad-spectrum star-polymer antiviral is to truly find use against a wide range of viruses, their efficacy in high protein environments (such as blood) needs to be explored and improved, especially for currently untreatable deadly viral infections such as dengue haemorrhagic fever.

Building on our previous findings, we synthesised star-polymers that incorporated varying percentages of a positively charged monomer, creating zwitterionic polymers, to prevent protein corona formation and enhance antiviral efficacy in high-protein environments. *In vitro* studies showed that 4-arm zwitterionic polymers with 12% positively charged monomer inclusion (ZP12), exhibited the most promising broad-spectrum virucidal properties in both low and high-protein environments against a wide range of viral families. This star-polymer was biocompatible *in vivo* when administered intraperitoneally (IP) and intravenously (IV). In a murine model of CHIKV infection, the zwitterionic polymer significantly reduced alphavirus-induced pathologies and the crosstalk between CHIKV immunopathogenesis drivers. We show that this star-polymer not only mitigates the negative effects of protein coronas, seen with our original star-PSS, but also maintains low toxicity and high antiviral efficacy *in vivo*, suggesting potential to combat future viral pandemics (Fig.1).

**Fig. 1:**
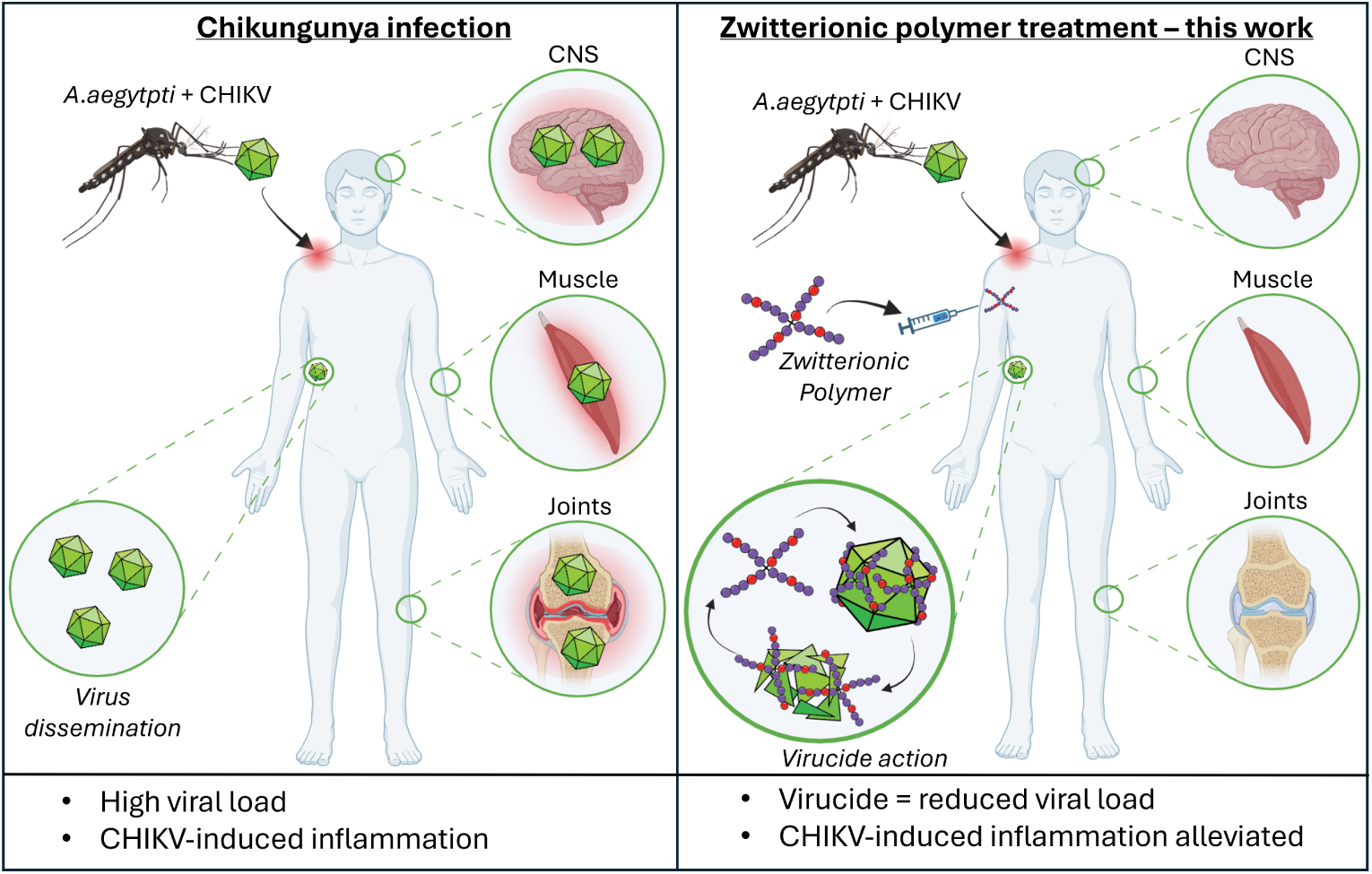
Schematic overview representing Chikungunya virus (CHIKV) infection and the associated pathologies (left panel) and suggested treatment approach using zwitterionic polymer treatment (right panel) showing reduced pathologies.

## Results and Discussion

### Zwitterionic polymer synthesis

Previously, our low toxicity star-PSS polymer was highly efficacious as a topical treatment for viral infections [45], however, in high protein biological fluids (such as blood), the highly negatively charged sulfonated moieties were shown to be coated by a protein corona, significantly reducing their antiviral efficacy[37, 56, 57]. This is problematic for viruses that disseminate in the blood (or other high protein environments)[61–64], such as CHIKV. It has previously been suggested that incorporating positively charged groups within sulfonated antivirals would help reduce or counteract the negative impact caused by protein corona’s[57]. Therefore, in order to allow for efficacy against CHIKV and other viruses in a high protein environment, we synthesised new star-zwitterionic polymers (ZPs) with varying percentages of the positively charged monomer, (vinylbenzyl)trimethylammonium chloride (PVB). We explored a range of PVB % inclusions, specifically 12% (ZP12), 47% (ZP47) and 97% (ZP97), alongside PSS (0%) (SI Fig.1-4). Full conversion was observed for each ZP, indicating that the target degree of polymerisation (DP) of ∼50 for each arm was reached (DP ∼200 total). The low polydispersity index (PDI) observed in the Dynamic Light Scattering (DLS) measurements indicate well defined star-polymers with similar sizes. However, variabilities in the hydrodynamic size and the zeta potential were observed. As PVB inclusion increased, hydrodynamic size decreased, suggesting a more compact polymer conformation, potentially due to charge interactions. No differences in zeta potential were observed between ZP12 and ZP47, however ZP97 showed a positively charged zeta potential, confirming a higher ratio of cationic monomer (SI Fig.1-4). Gel Permeation Chromatography (GPC) measurements indicate successful formation of the star-polymers, and the Mn values obtained for ZP12 and ZP47 further confirm that the molecular weight of these stars are of similar size. Results confirmed the successful synthesis of ZPs, which were consistent with the previously reported star-PSS (0% PVB inclusion)[45].

### Low toxicity and high efficacy in high protein environments

It was first necessary to confirm that the newly synthesised star-ZPs maintained the low toxicity and high antiviral efficacy profiles seen with the the parent star-PSS. To determine cell viability, the Cell Titre Glo assay, an ATP measurement for metabolically active cells, was used. ZP97 exhibited high toxicity (SI Fig.5), consistent with other positively charged polymers[65], and so was excluded from further testing. The remaining ZPs were all non-toxic *in vitro* even at the highest concentration tested (5000 *µ*g/mL), similar to star-PSS (SI Fig.6a-f). To probe the *in vitro* antiviral efficacies of the remaining ZP polymers, we infected HEK293T cells[11, 23, 66–70] with CHIKV IMT (MOI 1). The assay was conducted in both a low and high protein environment to determine if the inclusion of positive charge had reduced the negative impact on the antiviral efficacy due to the protein corona’s. In both instances, the media was supplemented with 2% fetal bovine serum (FBS), with the high protein environment having the addition of bovine serum albumin (BSA) (0.01%)[57]. A range of star-polymer concentrations, ranging from 51.2 pg/mL to 500 *µ*g/mL, were pre-incubated in each environment for 30 min at 37°C before incubating with CHIKV for 1 hr[45, 71]. The mixtures were then inoculated on to cell monolayers and infectivity assessed using antibody-labelling and flow cytometry, 24 hrs post infection (hpi).

In a low protein environment, star-PSS was the most effective antiviral (IC_50_ = 0.175 *µ*M, SI ∼ 41) (also observed against HSV-2, SI Fig.7) whilst ZP47 was the least effective (IC_50_ = 2.90 *µ*M, SI ∼ 2.4) (Table 1 and SI Fig.6g). This suggests that inclusion of positive charge (or less negative charge overall) reduces the antiviral efficacy of these star-polymers in low protein environments. However, the opposite effect is seen in a high protein environment (star-PSS IC_50_ = 2.65 *µ*M, and ZP47 IC_50_ = 0.81 *µ*M) (Table 1 and SI Fig.6h). These findings support the hypothesis that PVB inclusion leads to a reduced protein corona and enables the sulfonated moieties to bind viruses and exert an antiviral effect. For all further studies ZP12 is used, as it showed good efficacy in both environments, and uses the lowest % of positively charged monomer, which is known to have the least toxicity, to achieve improved antiviral properties in high protein environments[74].

**Table 1:** Inhibitory activities of star-polymers against CHIKV.

|  | Low protein environment |  |  | High protein environment |  |  | Cell viability |
| --- | --- | --- | --- | --- | --- | --- | --- |
| | IC<sub>50</sub><br>($\mu\text{g/mL}$) | IC<sub>50</sub><br>($\mu\text{M}$) | SI | IC<sub>50</sub><br>($\mu\text{g/mL}$) | IC<sub>50</sub><br>($\mu\text{M}$) | SI | CC<sub>50</sub><br>($\mu\text{g/mL}$) |
| PSS | 7.41 | 0.175 | >40.5 | 112 | 2.65 | >2.67 | >300 |
| ZP12 | 24.1 | 0.569 | >12.4 | 79.1 | 1.86 | >3.79 | >300 |
| ZP47 | 124 | 2.897 | >2.42 | 34.6 | 0.81 | >8.68 | >300 |

ZP12: 12%PVB. ZP47: 47%PVB. IC<sub>50</sub> inhibitory concentration at 50%. SI: selectivity index (CC<sub>50</sub>/IC<sub>50</sub>). CC<sub>50</sub>: cytotoxicity concentration at 50%. Based on previous toxicity studies involving antiviral polymers, SI values were calculated using 300 $\mu\text{g/mL}$ as the standard CC<sub>50</sub> value[29, 31, 39, 45, 71–73].

### Irreversible virucidal inhibitory mechanism

Sulfonated star-polymers demonstrated irreversible extracellular[45] *i.e.* virucidal mode of action (Fig.2a) and thus, we expect ZP12 to function similarly. To establish the mode of action, we performed time of addition assays[29, 71] (Fig.2b) against CHIKV. We pre-treated cells with various ZP12 concentrations (10 *µ*g/mL or 20 *µ*g/mL), followed by CHIKV infection to rule out intracellular effects. No inhibitory effects were observed, therefore excluding any intracellular cell-mediated effects. Next, we investigated the direct effects of ZP12 by adding it before, during or after CHIKV infection of cell monolayers. A directly proportional inhibitory effect was observed for each condition. The strongest inhibitory effect was observed with the 1 hr pre-incubation of CHIKV with ZP12, prior to inoculation onto HEK293T cells, elucidating that ZP12 needs direct contact with virions to exert its antiviral effects.

**Fig. 2:**
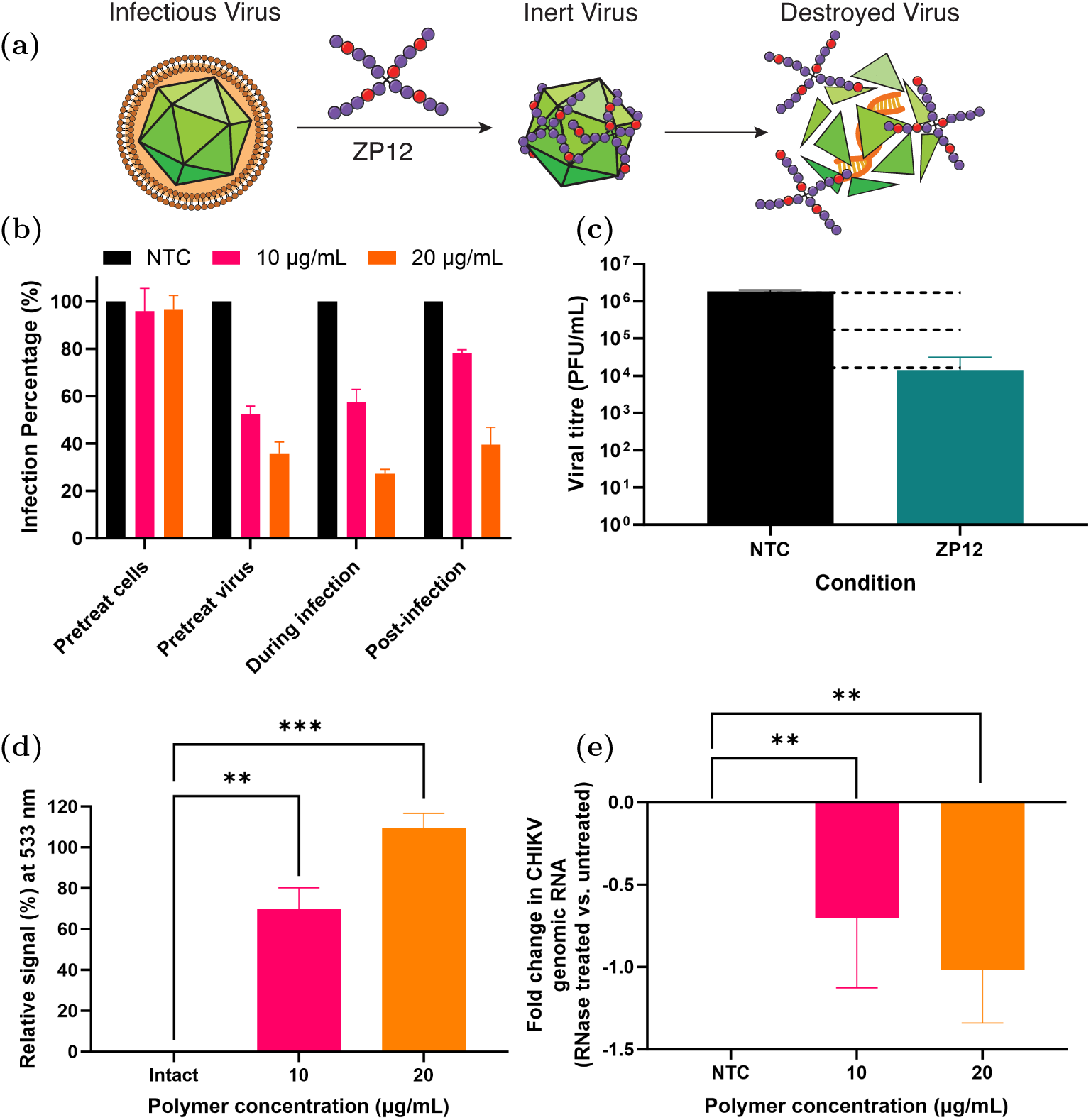
ZP12 mechanism of action. **(a)** Graphic illustration depicting ZP12 mechanism of action. **(b)** Time of addition assay suggests ZP12 only inhibits CHIKV when simultaneously added together and not before viral exposure. **(c)** Virucidal assay showing a 2-log reduction in CHIKV viral titre, indicating ZP12 (100 *µ*g/mL) has a virucidal mode of action. Viral titre was determined by comparing the treated and untreated wells. Dotted line represents 1-log and 2-log reductions, respectively. **(d)** FAIRY assay showing TO-PEG is accessible to CHIKV RNA following capsid destruction. Signals are normalised to 10 min 90^◦^C heated CHIKV sample. **(e)** RNA Exposure assay showing ZP12 (10 *µ*g/mL and 20 *µ*g/mL) is associated with the release of RNA from CHIKV, confirming the extracellular virucidal mechanism. Fold change difference (RNase - control) of CHIKV samples treated with or without ZP12. Results are mean*±*SEM from 3 independent experiments. Statistical analyses were performed using one-way ANOVA (^∗∗^p*<*0.01 and ^∗∗∗^p*<*0.001.)

To evaluate ZP12’s virucidal mode of action against CHIKV, we performed plaque-based virucidal assays (Fig.2c). The virucidal assay incubates ZP12 at its IC_90_ with CHIKV for 1 hr before infection of cell monolayers and subsequent serial dilution. The serial dilutions were performed to probe a virustatic mode of action[39], where if the polymer dissociates from the virus, infectivity would return. On the other hand, if the polymer exhibits a virucidal mode of action, a 2-log reduction in infectivity would be observed[45, 46, 71]. Here, we observed a 2-log reduction in infectivity when CHIKV was treated with ZP12, elucidating a virucidal mode of action[39, 45, 46, 71]. To further confirm a virucidal mechanism, we utilised the Fluorescence Assay for vIRal Integrity (FAIRY)[75] and an RNA exposure assay[51]. These assays evaluate the released RNA genome from damaged virions in a cell-free environment, providing an indication of viral infectivity. FAIRY uses thiazole orange terminated polyethylene glycol (TO-PEG), which is capable of binding to exposed nucleic acids, resulting in a fluorescence increase (normalised to heated sample with assumed 100% RNA release) when mode of action is virucidal. Whereas the RNA exposure assay utilises a RNase step to degrade free viral RNA, as a consequence of antiviral treatment, followed by qPCR quantification[31, 51, 71] with a lack of amplification indicating a virucidal mechanism. ZP12 showed a 70% increase in fluorescence, elucidating that the CHIKV virions were disrupted and consequently the viral RNA was exposed and bound to TO-PEG (Fig.2d). Additionally, a 0.7-fold decrease in viral RNA was observed (Fig.2e). These results further confirm the virucidal inhibitory mechanism of ZP12.

### ZP12 has broad-spectrum virucidal inhibitory properties

After confirming the extracellular virucidal properties of ZP12 against CHIKV, we investigated whether similar properties were observed for other CHIKV strains and different viral families (Fig.3a). We initially focused on exploring the effect of ZP12 against other CHIKV strains and a phylogenetically CHIKV-related alphavirus, namely O’nyong’nyong (ONNV). We incubated ZP12, in a dose response manner, with CHIKV Singapore (SG), Indian (IND) and Caribbean (CRB) strains; and ONNV IMT (Fig.3b). We found that irrespective of the strain, the IC_50_ values were ∼ 0.55 *µ*M (SI of ∼ 12.5), with the IC_50_ value of ONNV (IC_50_ = 0.58 *µ*M, SI ∼ 12.1) being similar. In parallel, virucidal assays confirmed the virucidal inhibitory properties of ZP12 against all CHIKV strains and ONNV (Fig.3c). The extracellular mode of action combined with the structural similarities of the virions of each strain, likely contribute to the similar IC_50_ results and virucidal mode of action, indicating that our approach is not strain or alphavirus specific.

**Fig. 3:**
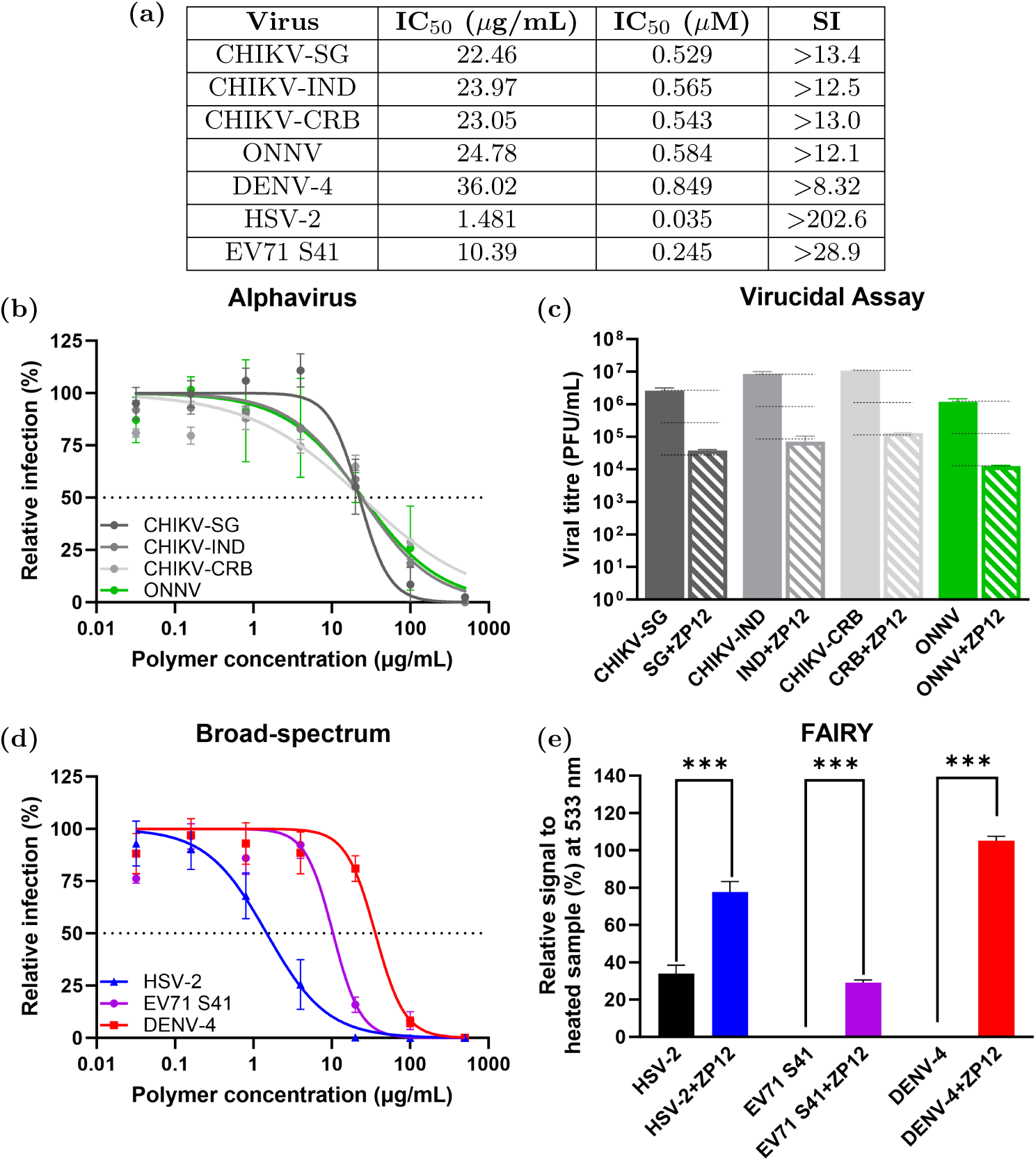
Broad-spectrum properties of ZP12. **(a)** IC_50_ and selectivity index (SI) summary. **(b,d)** Inhibitory effects against different CHIKV-strains (SG, CRB and IND), *Alphaviridae, Flaviviridae, Herpesviridae and Picornaviridae* families. Dotted line represents IC_50_. Percentage infections were expressed relative to signals from untreated infected cells. **(c,e)** Virucidal mechanism. **(c)** Virucidal assay showing 2-log reduction in viral titre (dotted line represents 1-log and 2-log reductions, respectively). Infection was determined by comparing the treated and untreated wells. **(e)** FAIRY showing fluorescence increase following capsid disruption. Signals are normalised to samples that underwent 10 minutes of 90^◦^C heating. Statistical analyses were performed using one-way ANOVA (^∗∗∗^p*<*0.001.) Data are presented as mean*±*SEM of three independent experiments performed in duplicate.

To further evaluate the broad-spectrum efficacy of ZP12, we investigated its antiviral properties against other viral families, specifically *Flaviviridae, Herpesviridae and Picornaviridae*. The Flaviviridae family is a group of viruses belonging to arboviruses, similar to alphaviruses. In particular we studied dengue virus serotype 4 (DENV-4) (MOI 1), and observed an inhibitory effect with an IC_50_ of 0.85 *µ*M (SI ∼ 8.3) (Fig.3d), with a confirmed virucidal mode of action (Fig.3e). Additionally, we observed a virucidal mode of action against two other flaviviruses, DENV-1, which is serologically similar to DENV-4, and Zika virus (SI Fig.8). It has been suggested that flaviviruses are not HSPG-dependent[9, 72], indicating that ZP12 may be effective against HSPG-dependent and HSPG-independent viruses, although strain adaptation to cell cultures needs to be considered[72].

To further probe the broad-spectrum efficacy of ZP12, we tested it against an enveloped DNA virus, HSV-2 (*Herpesviridae*); and a non-enveloped RNA virus, Enterovirus-71 (EV71) (*Picornaviridae*). The inhibitory properties were evaluated on Vero cells (Fig.3d) and the virucidal mechanism determined using FAIRY (Fig.3e). We observed an inhibitory effect for HSV-2 and EV71 with an IC_50_ of 0.035 *µ*M (SI ∼ 203) and 0.25 *µ*M (SI ∼ 28.9) (Fig.3a), respectively, with a confirmed virucidal mode of action (Fig.3e). Although the fluorescence obtained during the FAIRY measurements are higher for some viruses, this is due to the difference in viral titre and does not imply that ZP12 is a better virucide for certain viruses[75]. Overall, ZP12 demonstrated broad-spectrum virucidal inhibitory properties against a wide range of viral families, viral serotypes as well as enveloped and non-enveloped virions.

### CHIKV-induced joint swelling and tissue viral load is reduced

The *in vitro* biocompatibility and efficacy of ZP12 against a wide range of viruses is extremely promising, and so next we explored the *in vivo* properties using a C57BL/6 CHIKV murine model[69, 76, 77]. Compounds greater than 5 kDA, like ZP12 (42 kDA), administered intraperitoneally (IP) are known to drain into the lymphatic system[78]. Given that CHIKV disseminates through the lymphatic system before entering the bloodstream, we examined the biocompatibility of ZP12 in both blood (intravenous (IV) administration) and the lymphatic system (IP administration) (SI Fig.9a-c). Animals tolerated 10 mg/kg ZP12 via IP (used for further studies on account of it being less invasive) and 5 mg/kg ZP12 via IV every 24 hours for 7 days. We also investigated whether 5 mg/kg of the fully sulfonated star-PSS administered via IV (every 24 hours for 7 days) was tolerated by the animals. However, all animals died within 3 dpi (SI Fig.9d). The animals survival after being IV treated with ZP12 (5 mg/kg) highlights the significant improvement that the inclusion of 12% PVB has on the *in vivo* toxicity.

We then assessed the most effective regimen to reduce the CHIKV-induced inflammatory pathology (SI Fig.10a). To do so, 4-week-old C57BL/6 female mice were inoculated with 10^6^ CHIKV PFUs subcutaneously (s.c) on the ventral side of the right hind footpad. An hour post-inoculation (hpi), either 10 mg/kg ZP12 or the vehicle carrier (PBS) was administered IP as either i) 1 dose (1 hpi); ii) 3 doses (1 hpi, 2 days post infection - dpi, and 4 dpi); or iii) 7 doses (every 24 hours for 7 days). We hypothesised that ZP12 would reduce CHIKV virions, on account of its virucidal mode of action, and the subsequent CHIKV-induced pathologies. We monitored the CHIKV-induced joint inflammation at the site of infection, with the biphasic peaks being characteristic of CHIKV replication and the innate immune response (2-3 dpi); and infiltrating immune cells (6-7 dpi)[77]. Significant reductions were observed for both the 3 and 7 dose regimens, with a greater joint swelling reduction being observed with the 7 doses (every 24 hrs for 7 days) (Fig.4a-b and SI Fig.10b). Using the 7 dose regimen (every 24 hours for 7 days), we next evaluated whether a lower ZP12 dose remained effective. Using the same mouse model, we treated the animals with 1 mg/kg, 5 mg/kg and 10 mg/kg of ZP12 every 24 hours for 7 days (SI Fig.10c). We observed no difference in the peak associated with CHIKV replication and the innate immune response (2-3 dpi) with the 1 mg/kg and 5 mg/kg dosages, however a difference was observed at the second biphasic peaks with each dosage. However, the most significant reduction in joint swelling for both biphasic peaks was observed when the animals were treated with 10 mg/kg of ZP12 every 24 hours for 7 days (SI Fig.10b-e). The 7 dose (10 mg/kg every 24 hours for 7 days) regimen was used for the remainder of the studies.

**Fig. 4:**
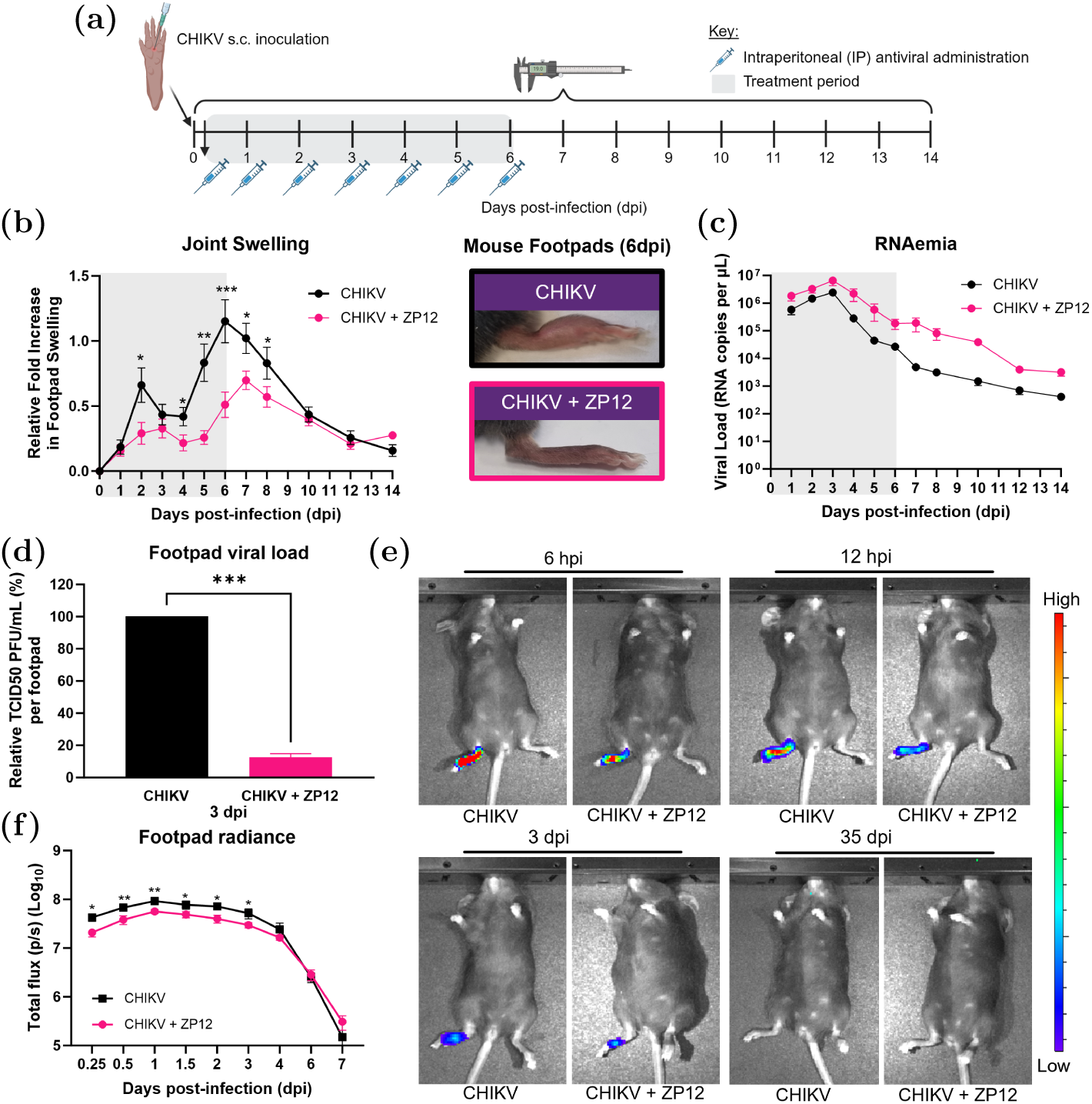
CHIKV tissue viral load and CHIKV-induced inflammatory pathologies are reduced when animals are intraperitoneally (IP) administered 10 mg/kg ZP12 every 24 hours for 7 days. **(a)** Schematic representing C57BL/6 animal infection with 10^6^ PFU CHIKV in the right footpad, were ZP12 treatment began 1 hour post-infection (hpi), as represented by the grey box. **(b)** Joint swelling with images at 6 dpi representing swelling decrease treated animals and **(c-d)** RNAemia and footpad viral load at 3 dpi in CHIKV only and CHIKV treated groups. **(e)** Representative pseudo-coloured images of bioluminescence readings showing localised CHIKV infection and reduction in viral load with ZP12 treatment at 6 hpi, 12 hpi, 3 dpi and 35 dpi. **(f)** Footpad radiance comparing CHIKV and CHIKV+ZP12 groups. Grey box represents treatment period. All data are presented as means±SEM of at least 5 animals per experimental group and are representative of two independent experiments (n=2). Comparison between treated and untreated groups were performed with non-parametric Mann-Whitney U test (two-tailed ^∗^p*<*0.05, ^∗∗^p*<*0.01 and ^∗∗∗^p*<*0.001.)

Next, we investigated the impact ZP12 had on other CHIKV-induced pathologies, including RNAemia. Typically, qPCR is used to investigate RNAemia, with an expected decrease in viral RNA being associated with successful treatment. However, due to ZP12’s virucidal mode of action, the released RNA would still be detected by qPCR, thus the values obtained would correlate to the combination of RNA from both infectious and non-infectious virions. Thus, the use of qPCR to determine viral load when investigating virucidal antivirals is not appropriate. As expected, we observed no significant difference in RNAemia over the course of the study (Fig.4c) CHIKV inflammation is localised to the site of infection and in order to more accurately determine the viral load post-treatment, we harvested the infected joint at the peak of RNAemia infection (3 dpi) enabling the accurate assessment of the infectious CHIKV titre using plaque assays. Here, we observe a significant reduction in the tissue viral load of *>*90% (Fig.4d), achieved using IP administration elsewhere from the site of infection.

To assess any possible differences in CHIKV replication kinetics at the site of inoculation (right hind limb footpad), viral replication *in vivo* was assessed using a luciferase-tagged CHIKV clone that mimics wild-type CHIKV infection in mice[79]. The replication cycle of CHIKV is proportional to the luciferase activity, meaning as CHIKV replicates, the viral RNA is translated into viral proteins and a luciferase enzyme. To quantify the luciferase enzyme produced, the animals were exposed to a luciferin substrate, resulting in the emittance of bioluminescence light. The bioluminescence readouts correlate with the viral burden found in infected footpads. A significant difference in footpad radiance was detected as early as 6 hpi in ZP12 treated mice, the time in which CHIKVs first replication cycle occurs[80, 81](Fig.4e-f). The bioluminescence measurements indicated that ZP12 not only effectively reaches the infection site (when administered elsewhere) but also reduces viral replication within 5 hours post administration (hpa). The reduction in bioluminescence signals were maintained until it dissipated by 4 dpi. The reduction in tissue viral load and subsequently the bioluminescence signals demonstrate that ZP12 is effective at reducing CHIKV replication and the subsequent viral progeny produced are also destroyed upon contact with ZP12. More importantly, a longitudinal study confirmed that the bioluminescence signal did not return up to 35 dpi (Fig.4e).

To determine the broad-spectrum efficacy of ZP12 *in vivo*, we also infected 4-week-old C57BL/6 female mice with 10^6^ ONNV IMT PFUs on the ventral side of the right hind footpad. Unlike CHIKV, ONNV provokes an immune response skewed towards the Th2 profile, which dampens viral inflammation and reduces tissue damage[82]. This model ensures that ZP12’s effectiveness can be evaluated across viruses with varying immunopathological responses. Following ONNV infection, each mouse was treated using the same regimen as the CHIKV studies (10 mg/kg every 24 hours for 7 days via IP administration). Similar to the CHIKV studies, we observe significant reductions in both joint swelling at 6 dpi and in tissue viral load at 3 dpi (SI Fig.11). These findings demonstrate that ZP12 is effective at reducing tissue viral load and inflammation following IP administration, highlighting its potential as a treatment for not only CHIKV and ONNV infections, but also other viremia-producing viruses such as Rift Valley Fever virus and Ebola virus.

### Reduction of cellular infiltration

Previous studies have shown that CHIKV-induced inflammation is exacerbated by the infiltration of leukocytes into the infected joint-footpad[5, 68, 83, 84]. Therefore, to assess the immune cells responsible for the joint swelling, we harvested the infected joint-footpad and the closest draining lymph node i.e. the popliteal lymph node (pLN). The pLN is the site in which immune cells expand and proliferate, before infiltrating the infected footpad joints. After isolating the cells from these tissues, the immune subsets were profiled using an antibody panel and flow cytometry[5]. Data obtained were visualised with the dimensional reduction tool, UMAP (Uniform Manifold Approximation and Projection)[85], normalised to 5,000 cells per joint-footpad and represent the proportion, but not the quantity, of cells present in these joints.

A significant increase in Ly6G^+^ neutrophils was observed in the UMAP analysis (Fig.5a) for animals dosed with ZP12 (CHIKV-infected and uninfected), suggesting some inflammatory response is being triggered by ZP12. To understand the potential ZP12 effect on the neutrophils, we profiled the circulating neutrophils (in blood) using an antibody panel and flow cytometry (SI Fig.12a). It was observed that the animals dosed with ZP12, had an increase of circulating neutrophils up to 7 dpi, where hereafter it began decreasing. This gave the notion that ZP12 was either accumulating in organs such as the liver or kidneys[86, 87], or degrading into toxic metabolites. To determine if these organs were being damaged by ZP12, we assessed the serum changes, of liver and kidney enzymes, which indicated no potential organ damage (SI Fig.12b). Therefore, it can be elucidated that the increase of circulating neutrophils is likely the result of ZP12 being present, and will not be further studied. UMAP analysis revealed substantial differences for the remaining immune cell populations, between the CHIKV-infected (treated and non-treated groups), with no substantial differences being observed between the control groups (carrier and ZP12 only) (Fig.5a). To further understand these difference, the immune cell subsets, based on the surface expression of the lineage markers, was plotted (Fig.5b and SI Fig.13). Residential dendritic cells (DCs) in the footpad are considered key initiators for CHIKV immunopathogenesis. DCs capture, process and present CHIKV antigens to other immune cells after migrating to the draining pLN, where immune cell activation and proliferation occur. In contrast to the control groups (ZP12-only and carrier), a significant reduction in DCs within the footpad was observed at 6 dpi in both CHIKV-infected groups (treated and non-treated). This reduction was anticipated, as the presence of high viral titres in the footpad, would drive the maturation and migration of DCs to the pLN, therefore reducing DCs in the footpad. We previously demonstrated that ZP12 was effective at reducing CHIKV viral load at the infected joint-footpad (Fig.4d). Based on this reduced tissue viral load and consequently viral antigens, less DCs mature in the footpad and drain into the pLN (SI Fig.13b). Dendritic cells are crucial antigen presenting cells that prime and activate CD4^+^ T cells. Previous studies showed that the pathogenesis of CHIKV is driven by the infiltration of CD4^+^ T cells[79] and the subsequent crosstalk with CD64^+^MHCII^+^ macrophages[5]. Gating on CD45^+^CD3^+^CD4^+^ revealed that the expression of CD4^+^ T cells in the footpad was downregulated following ZP12 treatment of CHIKV-infected mice (Fig.5b). In conjunction, a downregulation of CD11b^+^Ly6C^+^ infiltrating monocytes and consequently CD64^+^MHCII^+^ macrophages was observed. Ultimately, this reduction in activated immune cells results in less oedema in the infected joint-footpad, explaining the observed reduction in joint inflammation (Fig.4b). CD8^+^ T cells and NK cells, whose roles in the immunopathogenesis of CHIKV is not fully understood, were also reduced following ZP12 treatment of CHIKV-infected animals[84, 88].

**Fig. 5:**
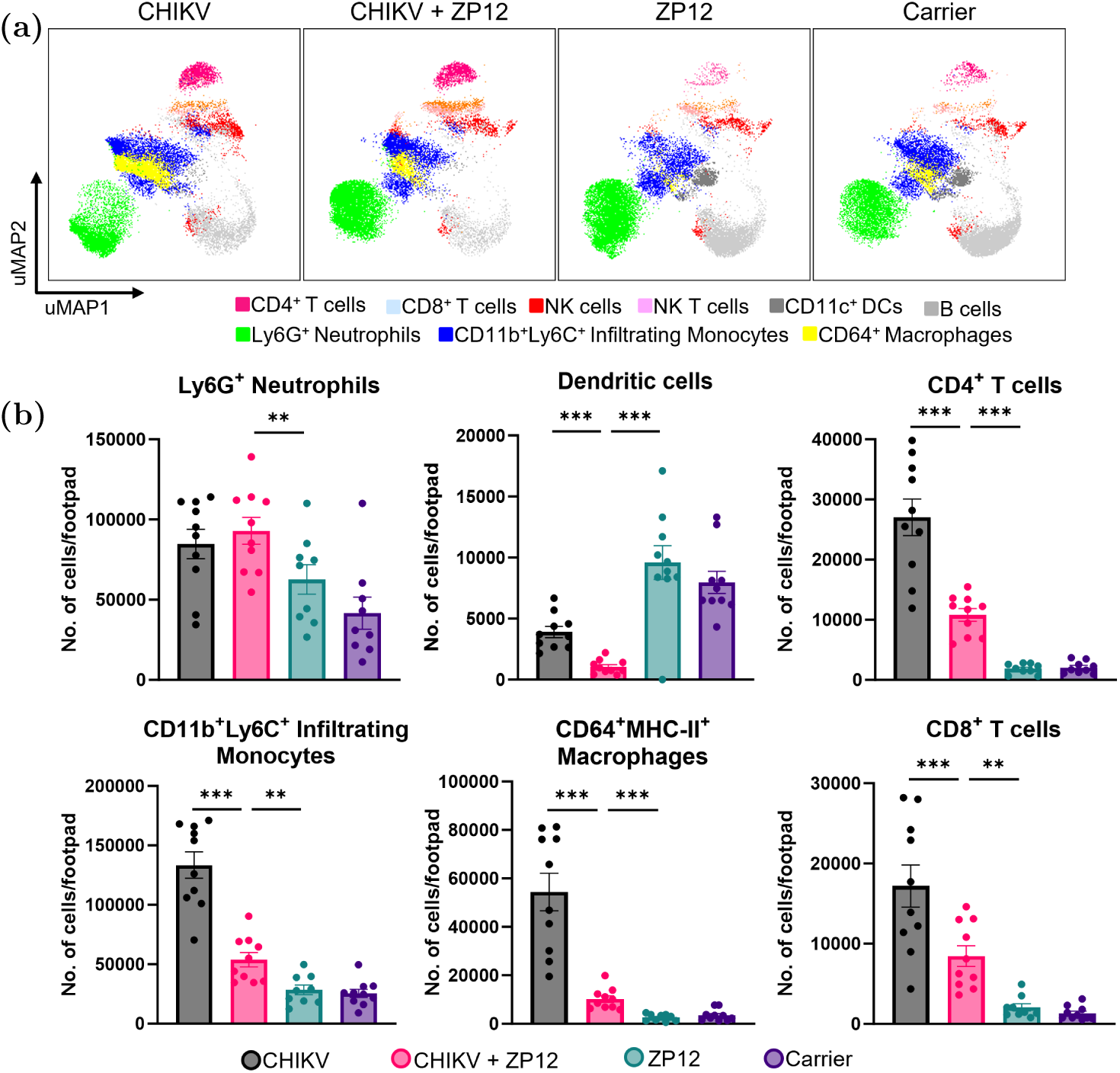
High-dimensional analysis of flow cytometry data reveals a decreased joint immune profile of infiltrating immune cells of CHIKV-ZP12 treated animals. **(a)** Joint-footpad cells were harvested at 6 dpi and stained with a panel of antibodies targeting lineage cell surface marker expression. Data were analysed by downsizing and concatenation of samples containing 8,000 randomly selected live CD45^+^ cells from each sample (n=5 per group) enabling the generation of a Uniform Manifold Approximation and Projection (UMAP). **(b)** Leukocytes responsible for CHIKV joint inflammation including CD4^+^ T cells, CD8^+^ T cells, NK cells, neutrophils, infiltrating monocytes and CD64^+^ macrophages were quantified at 6 dpi. Data are presented as mean±SEM of at least 5 animals per experimental group and are representative of two independent experiments (n=2). Comparison between treated and untreated groups were performed with non-parametric Mann-Whitney U test (two-tailed; ^∗∗^p*<*0.01 and ^∗∗∗^p*<*0.001).

Studies have shown that the immunopathogenesis of alphaviruses is mediated by specific T helper (Th) cells[83, 89, 90]. For example, CHIKV inflammation is mediated by Th1 cells that produce pro-inflammatory cytokines including IFN-*γ* and TNF-*α*, whilst ONNV is mediated by Th2 cells that have anti-inflammatory properties due to the release of cytokines such as IL-4. Given the importance of CD4^+^ T cells in the pathology of CHIKV infections, we needed to ensure that the reduction in CD4^+^ T cells of ZP12 treated CHIKV-infected animals was not due to ZP12 being immunosuppressive or altering the immune phenotype of the Th1 cells. To investigate this, we incubated various ZP12 concentrations on CD4^+^ T cells isolated from naive spleens. To quantify the cytokines produced by CD4^+^ T cells, we stimulated the cells with T cell-activating agents phorbol myristate acetate (PMA) and ionomycin. Using an intracellular antibody-labelling panel and flow cytometry (SI Fig.14a), no differences in any cytokine production was observed, elucidating that ZP12 was not altering the phenotype of CD4^+^ T cells or the mediators produced (SI Fig.14b). Studies have shown that the crosstalk between immune subsets drives the immunopathogenesis of CHIKV infections[5, 84, 91], we aimed to investigate the effect any mediators (chemokines and cytokines) would produce in the presence of a whole immune environment. To assess this, we exposed splenocytes to various ZP12 concentrations overnight, followed by either T-cell stimulation with PMA/ionomycin, or a macrophage activating agent, lipopolysaccharide (LPS) (SI Fig.14c). Mediators were subsequently quantified using a fluorescent bead-based multiplex immunoassay and plotted visually on a heat map. Irrespective of the macrophage or T cell mediators measured, no differences were observed.

After confirming that ZP12 did not alter the phenotype or mediators produced by T cells and macrophages, we decided to investigate whether ZP12 would inhibit these mediators *in vivo*. If ZP12 was truly suppressing the immune response, we would not have observed any reductions in joint swelling when animals were treated with ZP12 after the onset of the disease (SI Fig.14d-e). To study this, we infected mice with CHIKV as previously described, and began ZP12 treatment (10 mg/kg every 24 hours for 4 days via IP) after the known peak of viremia i.e. 3 dpi. As expected from the *in vitro* data, no significant difference in joint swelling was observed, further confirming that ZP12 was not suppressing the immune system.

## Conclusion

Arboviruses, including CHIKV, have potential to cause the next pandemic linked with autochthonous transmission. To date no effective antiviral therapeutics exist. Our group previously synthesised a low toxicity and highly efficacious sulfonated star-polymer (star-PSS) with topical therapeutic properties, but the antiviral efficacy decreased in biological fluids as a result of protein corona formation. Therefore, on the basis of this work, we synthesised star-polymers with inclusion of varying percentages of a positively charged monomer, PVB. We anticipated the addition of positive charge would reduce protein corona formation, and consequently increase the antiviral efficacy.

ZP12 demonstrated the most promising broad-spectrum inhibitory virucidal properties in both low and high protein environments, against a wide range of viral families including *Alphaviridae, Flaviviridae, Herpesviridae, and Picornaviridae*; viral serotypes; as well as enveloped and non-enveloped viruses *in vitro*. We further probed whether ZP12 retained these antiviral properties in high protein biological fluids using a C57BL/6 CHIKV murine model. ZP12 was biocompatible up to 10 mg/kg via IP administration and 5 mg/kg via IV administration every 24 hours for 7 days. More importantly, ZP12 was capable of significantly alleviating CHIKV-induced joint swelling, through the reduction of viral replication within 5 hpa of ZP12, as well as significant reduce tissue viral load by 90%. Similar results were also obtained for a phylogenetically CHIKV-related alphavirus, namely ONNV. Consistent with the reduced viral burden, fewer immune cell subsets known to exacerbate CHIKV-induced inflammation were activated. Specifically, we identified a significant reduction in the crosstalk between the recruitment of CD4^+^ T cells and the subsequent activation of CD64^+^MHCII^+^ macrophages. Collectively these findings established ZP12 as a strong therapeutic candidate against alphaviruses.

Given the documented impact of age and sex on viral disease progression and immune response, future work should focus on expanding these findings to encompass a range of biological contexts (*i.e.* gender and age) to ensure the generalisability and translational relevance. Moreover, the efficacy of ZP12 has thus far only been validated *in vivo* against alphaviruses. To fully explore its therapeutic potential, further studies are required to assess its activity against a broader spectrum of viral infections, particularly against more severe and life-threatening systemic viral infections such as DENV, Zika virus and Ebola virus.

In summary, this study has suggested that ZP12 is not only capable of being effective in high protein environments but can also be administered elsewhere from the site of infection, circulate systemically and reach virions (<u>without</u> virus-specific targeting mechanisms antibodies). Importantly, this work provides the first evidence that broad-spectrum, extracellular, star-shaped ZP antivirals are a highly effective and promising approach for treating systemic viral infections. As such, ZP12 has been positioned as an extremely promising antiviral intervention with significant potential to help counter future viral outbreaks and enhance pandemic preparedness. As expected, due to reduced viral load, we also show that less immune cell subsets known to exacerbate CHIKV-induced inflammation were activated, resulting in a significant decrease in the crosstalk between the recruitment of CD4^+^ T cells and subsequent CD64^+^MHCII^+^ macrophages activation.

## Materials and Methods

Reagents were purchased from ThermoFisher Scientific unless otherwise stated. All reagents were handled following sterile techniques in biological safety cabinets.

### Cell lines

Human embryonic kidney HEK293T cells (CRL-11268, ATCC), and African green monkey kidney Vero-E6 cells (CCL-81, ATCC) were propagated in Dulbecco’s modified Eagle medium (DMEM) supplemented with 10% heat-inactivated fetal bovine serum (FBS, Gibco) and 1% Penicillin-Streptomycin (P/S, Gibco) (hereafter referred to as medium). Cells were mycoplasma free and maintained at 37^◦^C with 5% CO_2_. A mosquito cell line *Aedes albopictus* C6/36 cells (CRL-1660, ATCC), were propagated in Leibovitz’s L-15 medium (Life Technologies), supplemented with 10% FBS. Cells were maintained at 28^◦^C in a CO_2_ free environment.

### Viruses

CHIKV isolates: (i) IMT (LR2006-OPY1) was isolated from a French patient returning from the La Reunion Islands during the 2006 outbreak[92, 93]; (ii) SG (SGP011) was isolated by the National University Hospital following an outbreak in Singapore in 2008[93–95]; (iii) IND was isolated from the cerebrospinal fluid of an Indian patient during the 2010 outbreak (kindly donated by Dr. Ooi Eng Eong, DUKE-NUS Graduate Medical School, Singapore)[96]; and (iv) CRB (CNR20235) was isolated from a patient during the Caribbean island outbreak in 2013[89]. The ONNV-IMT (SSA/5163) isolate was isolated from an infected patient from Chad in 2004 (kindly donated by Marc Grandadam from Unite de Virologie Tropicale, IMTSSA, Marseille, France)[82, 92, 97]. HSV-2 samples were originally isolated, verified and kindly donated by the University of Manchester, School of Medical Sciences (Professor Pamela Vallely). DENV-2 and DENV-4 were isolated from patients in Singapore (kindly provided by the National Public Health Laboratory, Singapore). EV71 S41 (5865/SIN/00009) was isolated during the Singapore outbreak in 2000[98]. The CHIKV variant expressing firefly luciferase (CHIKV-Fluc) was generated from full-length infectious cDNA clones of CHIKV-IMT[68, 79, 83, 94]. For *in vitro* studies, viruses were propagated and quantified by standard plaque assay in Vero-E6 cultures. For *in vivo* studies, viruses were propagated in *Aedes albopictus* C6/36 cultures, purified by sucrose-gradient ultracentrifugation[99] and quantified by standard plaque assay in Vero-E6 cultures.

### Cell Cytotoxicity Assays

Cell viability was assessed using the CellTiter-Glo luminescent assay (Luciferin, Promega) to measure ATP levels, as previously described[100]. Briefly, HEK293T and Vero-E6 cells were seeded (1.5E^4^ cells/well) in 96-well plates and incubated overnight. Cells were treated with serial dilutions of PSS, ZP12, ZP47 and ZP97 (50 to 5000 *µ*g/mL) in triplicates. Untreated controls contained cells in medium, while blank controls contained medium only. After 24 hrs, 100 *µ*L of CellTiter-Glo reagent was added as per the manufacturer’s protocol. Luminescence was measured after a 5 min incubation using the GloMax Multi-detection system (Glomax Explorer Software, v4.0, Promega). The 50% cytotoxic concentrations (CC_50_) were determined using Prism software (Graph-Pad Software, San Diego, CA).

### Viral Inhibition Assay

HEK293T cells or Vero-E6 cells were seeded (8.5E^4^ cells/well) in 24-well plates overnight. PSS, ZP12 and ZP47 were serially diluted (51.2 pg/mL to 500 *µ*g/mL) in medium and incubated with virus at a multiplicity of infection (MOI) of 1, for 1 hr at 37^◦^C. Cell medium was replaced with the virus-polymer mixtures allowing virus adsorption (1 hr at 37^◦^C). Subsequently, the virus inoculum was removed and cells were incubated in MTC medium (3:7 ratio of 1.5wt% methyl cellulose in deionised water and 2% FBS, 1% P/S DMEM) until cytopathic effects (CPE) were observed. Infection was assessed by either i) 0.5% crystal violet and plaque counting (HSV-2), or ii) antibody-labelling and flow cytometry analysis (CHIKV strains, ONNV, EV71 and DENV-4). The concentration of the compound producing 50% reduction in virus infection (IC_50_) was determined using the Prism software by normalising the percentage of infection for the drug-treated and untreated wells.

For CHIKV antibody-labelling, live cells were stained with a Live/Dead Fixable Aqua dye (Invitrogen) for 15 min, fixed and permeabilised (BD Sciences). Subsequently for CHIKV infection, cells were incubated with rabbit anti-CHIKV nsP2 [HL1431] primary antibody (GTX636897, GeneTex) (1:4000) for 10 min, followed by the secondary antibody, Alexa Fluor 488-conjugated Goat anti-rabbit IgG (1:2000) for a further 10 min. For acquisition, cells were resuspended in PBS and acquired using the NovoCyte Penteon (NovoExpress software, v4.0.5). Similar antibody-labelling methods (with different antibodies) were used for ONNV, EV71, and DENV-4. For ONNV, rabbit anti-ONNV nsp2 primary antibody (1:2000) and Alexa Fluor 488-conjugated Goat anti-rabbit secondary IgG (1:2000) was incubated for 10 min each. For EV71, mouse anti-EV71 primary antibody (1:2000) (MAB979, Merck) and Alexa Fluor 488-conjugated Goat anti-mouse secondary IgG (1:500) was incubated for 10 min each. For DENV-4, human anti-dengue primary antibody (hu1B, 1:6000) and Alexa Fluor 647-conjugated Goat anti-human secondary IgG (1:500) was incubated for 1 hr each.

### Protein Adsorption Viral Inhibition Assay

Same method as above, except PSS, ZP12 and ZP47 were serially diluted and incubated in medium containing 0.1 mg/mL sterile Bovine Serum Albumin solution (BSA) (0.01%) for 30 min at 37^◦^C[57], before being incubated with CHIKV or HAV-2 for 1 hr.

### Time of Addition Assay

ZP12 (10 or 20 *µ*g/mL) was incubated on HEK293T cells 1 hr before, during, or after CHIKV infection (MOI 1)[29, 71, 101]. Viral titres were then quantified using antibody-labelling and flow cytometry as described above, with results normalised to the non-treated control (NTC).

### Virucidal Mechanism: Virucidal Assay

The virucidal assay, assessed by TCID_50_, determines the polymers mode of action[29, 71]. Briefly, a 1:1 ratio of CHIKV and ZP12 (IC_90_, 100 *µ*g/mL), were incubated for 1 hr at 37^◦^C. The NTC was treated with PBS. For each condition, three Eppendorf tubes with 450 *µ*L medium were prepared. After incubation, 50 *µ*L of CHIKV-ZP12 and CHIKV-PBS mixtures were transferred to the first Eppendorf tube and serially diluted into the remaining two tubes (1:30, 1:300, 1:3000). 50 *µ*L from each dilution was added to confluent Vero-E6 cells (1.5E^4^ cells/well) in wells A1-A4, A5-A8, and A9-A12, respectively, followed by a serial dilution down the plate. Afterwards, incubate until visible CPE formation, cells were stained with 0.5% crystal violet. Viral infectivity was assessed using light microscopy and viral titres were quantified using the Reed-Muench method and converted to PFU/mL.

### Virucidal Mechanism: FAIRY assay

FAIRY serves as a nucleic acid fluorescent readout for viral infectivity[75]. Briefly, ZP12 and virus in Phenol Red-free DMEM (with no FBS supplementation) were incubated at a 1:1 ratio for 1 hr at 37 ^◦^C. The NTC was incubated with PBS, the blank controls were medium only, and the positive control, was virus heated at 90 ^◦^C for 10 min. Following incubation, 50 µL of TO-PEG and 50 µL of sample was added to a black 96-well flat bottom plate and incubated for 10 min. Fluorescent readouts were measured using the Tecan Microplate Reader Infinite 200 Pro system (Infinite RX software, v5.31) and normalised to the heated control.

### Virucidal Mechanism: RNA Exposure assay

RNA Exposure Assay detects released viral genomic RNA by quantitative RT-PCR (qRT-PCR) upon the addition of the virucidal compound[29, 51]. Briefly, a 1:1 ratio of CHIKV (1E^5^ pfu/mL) and the desired ZP12 concentration was incubated for 1 hr at 37 ^◦^C, followed by a 1:20 PBS dilution. 1.1 *µ*L 10 mg/mL RNAse A (T3018-2, BioLabs) or Milli-Q water was added into 10 *µ*L PBS buffer, followed by 100 *µ*L of the 1:20 diluted viral solution.

After 30 min incubation at 37^◦^C, RNA was extracted using the QIAamp Viral RNA kit (QIAGEN), following the manufacturer’s instructions. Samples were quantified by qPCR using a QuantiTect Probe RT-PCR Kit (QIAGEN), CHIKV E1 RNA standards and the Bio-Rad CFX Opus 384 RT-qPCR machine (Bio-Rad)[89]. Fold change was calculated using the ΔCt method.

### Mice

Four-week old female C57BL/6J mice were used for all experiments, unless otherwise stated. Animals were bred and kept under specific pathogen-free conditions in the Biological Resource Centre (BRC), Agency for Science, Technology and Research (A^∗^STAR), Singapore. All experimental procedures were approved by the Institutional Animal Care and Use Committee (IACUC: 211635) of A^∗^STAR, in accordance with the guidelines of the Agri-Food and Veterinary Authority and the National Advisory Committee for Laboratory Animal Research of Singapore.

### Biocompatibility of ZP12 *in vivo*

ZP12 (100 *µ*L) was administered via an intraperitoneal (IP) or intravenous (IV) route at 33 mg/kg, 10 mg/kg and 5 mg/kg every 24 hrs for 7 days. Four-week old male C57BL/6J mice were used for the *in vivo* intravenous (IV) PSS and ZP12 protein corona studies. PSS (100 *µ*L) was IV administered at 5 mg/kg and 1 mg/kg every 24 hours for 7 days. Animals bodyweight and behavioural changes using the grimace scale (score of 0=no pain, 1=moderate pain, 3=obvious pain)[102] were monitored daily up to 8 days post-administration (dpa) and then on alternate days until 14 dpa. If animals showed obvious signs of pain with significant reductions in bodyweight (*>*20%), they were humanely euthanised.

### Virus infection and evaluation of disease

CHIKV infection in mice were performed by inoculating 10^6^ PFUs CHIKV (in 30 *µ*L PBS) subcutaneously in the ventral side of the right hind footpad near the ankle. Following 1 hr post-infection, 100 *µ*L of 10 mg/kg ZP12 was administered via IP every 24 hrs for 7 days (for the therapeutic study, ZP12 was administered from 3 dpi up to 6 dpi). For effectiveness of the star-polymers in the blood, 1 mg/kg and 5 mg/kg ZP12 and PSS were IV administered every 24 hrs for 7 days. For the non-treated groups, 100 *µ*L PBS was administered. Joint swelling at the footpad was scored daily, starting from 0 day post-infection (dpi) until 8 dpi, and then on alternate days until 14 dpi. Measurements were made for both the height (thickness) and the breadth of the metatarsal region of the footpad and were quantified as (height × breadth) using electronic callipers. The degree of swelling was expressed as the relative increase in footpad size compared with the same footpad pre-infection (day 0), using the following formula: (*x −* day 0)*/*day 0, where *x* is the quantified footpad measurement for each respective day. Viremia was monitored daily from 1 to 8 dpi and then on alternate days until 14 dpi. Briefly, 10 *µ*L of tail blood from each mouse was collected and mixed in 120 *µ*L of PBS supplemented with 10 *µ*L of citrate-phosphate-dextrose solution (citrate). Viral RNA extractions were performed using the QIAamp Viral RNA kit (QIAGEN) following the manufacturer’s instructions. Viral copies were quantified by qRT-PCR using a QuantiTect Probe RT-PCR Kit (QIAGEN) and the Bio-Rad CFX Opus 384 RT-qPCR machine (Bio-Rad) with previously described methods that detect for CHIKV nsp2[76, 79, 96].

### Tissue viral load determination

Virus-infected footpads were harvested at 3 dpi, shredded in 2% complete DMEM and vortexed for 15s at 4000rpm (repeat twice) on the GentleMACS Dissociator using the GentleMACS M-tubes with strainer (Miltenyibiotec). The supernatant was filtered through a 40 *µ*m cell strainer. Infectious titre was assessed using TCID_50_. Briefly, 50 *µ*L of the supernatant was added to the cells, and serial dilutions were performed. Plates were incubated for 48 hours at 37^◦^C, until CPE was visible. Cells were stained with 0.5% crystal violet solution. Viral infectivity was assessed by cytopathic effects observed in wells via light microscopy. Viral titres were quantified using the Reed-Muench method and converted to PFU/mL.

### *In vivo* viral replication assay

To monitor virus replication, mice were inoculated with a firefly luciferase-tagged CHIKV infectious clone (CHIKV-Fluc), and tracked using the IVIS Spectrum In Vivo Imaging System (Perkin Elmer, Lumina S5). Bioluminescence signals were assessed daily from 1 to 8 dpi, then weekly up to 35 dpi. Animals were anaesthetised throughout the experiment in an oxygen chamber with 2% isoflurane. Mice were subcutaneously injected with 100 *µ*L of the luciferase substrate, D-luciferin potassium salt (Perkin Elmer), which was prepared by dissolving in PBS at a concentration of 10 mg/mL. Bioluminescence readings were taken 5 minutes post D-luciferin injection with a field of view (FOV) of 21 cm with exposure time for 20 seconds. Bioluminescence signals of the region of interest were drawn using the software Living Image (Perkin Elmer v 4.7.4) and total flux values (photons/second) were calculated.

### Popliteal lymph node and footpad leukocyte profiling

The popliteal lymph nodes (pLN), and right footpads of treated and non-treated animals, were harvested at 6 dpi as previously described[79, 82, 89, 93, 103]. Briefly, the harvested tissues were digested in digestion medium containing Dispase II (2 U/mL; Roche Applied Science), Collagenase IV (20 *µ*g/mL; Sigma-Aldrich) and DNase I (50 *µ*g/mL; Sigma-Aldrich) in complete RMPI medium (referred to as digestion medium) and agitated slightly at 37 ^◦^C (pLN: 30 min and footpad: 3 hrs). The tissues were filtered through a 70 *µ*m nylon mesh cloth (Sefar) and 40 *µ*m cell strainer (Fisherbrand), respectively. Red blood cells were lysed with 1×Flow Cytometry Mouse Lysis Buffer (R&D Systems) and footpad cells were purified with 35% v/v Percoll (Sigma-Aldrich) in FBS/RPMI. Isolated cells were enumerated and stained with Live/Dead Fixable Aqua dye (Invitrogen). Cells were washed and resuspended in 30 *µ*L of blocking buffer containing TruStain FcX PLUS antibody (anti-mouse CD16/32, clone S17011E, Biolegend)(1:200) diluted in PBS and incubated in the dark for 20 min. pLN cells were stained with the following antibodies: BUV395-conjugated anti-CD45 (clone 30-F11), PE-Cy7–conjugated anti-CD3 (clone 17A2), PE-conjugated anti-CD4 (clone GK1.5), PB-conjugated anti-CD8 (clone 53-6.7), PB-conjugated anti-B220 (clone 53-6.7), BV650-conjugated anti-CD11b (clone M1/70), Qdot605-conjugated anti-CD11c (clone N418), APCCy7-conjugated anti-Ly6C (clone HK1.4), AF700-conjugated anti-MHC-II (clone M5/114.15.2) and PerCP5.5-conjugated anti-LFA-1 (clone H155-78) for 30 min on ice. The same antibody panel was utilised for the footpad cells with the addition of Biotin-conjugated anti-NK1.1 (clone PK136), BUV737-conjugated Streptavidin, PE-CF594-conjugated anti-Ly6G (clone IA8) and APC-conjugated anti-CD64 (clone X54-5/7.1). All antibodies used for staining were purchased from BD Biosciences, BioLegend or ThermoFisher Scientific. Stained cells were fixed with IC Fixation Buffer (Invitrogen) and acquired using the BD FACSymphony A5 (BD FACS Diva software v8.0; BD BioScience) and analysed with FlowJo v10.6.2 (Becton, Dickinson and Company), using previously defined gating strategy[5].

### High dimensional analysis of flow cytometry data

Live CD45^+^ singlet events were pre-gated and randomly downsampled to a fixed number (n = 5,000) for each sample using FlowJo v10.6.2. Down-sampled files were concatenated and analysed using UMAP for Dimension Reduction plug-in v3.1 using default parameters (number of nearest neighbours = 15, minimum distance = 0.1).

### Statistical analysis

Animals were randomised prior to starting each experiment. Comparisons between the different experimental groups or samples were performed using non-parametric two-tailed Mann-Whitney test. Data were presented as mean *±* SEM, unless otherwise stated. Statistical analyses were performed using GraphPad Prism version 9.1. P-values considered statistically significant are represented with ^∗^p*<*0.05, ^∗∗^p*<*0.01 and ^∗∗∗^p*<*0.001.

### Star polymer characterisation

^1^H NMR (nuclear magnetic resonance) (400 MHz) spectra were recorded using a Bruker Avance III. Chemical shifts were recorded in parts per million (ppm) (*δ*) in deuterium oxide (*D*_2_O) (4.79 ppm). Aqueous gel permeation chromatography (GPC) measurements were carried out using a GPC eluent containing phosphate buffer (pH 9.0, ambient temperature) with 30% v/v methanol at a flow rate of 1.0 mL/min. The 1260 Infinity II Multi-detector GPC was fitted with two PL aquagel-OH MIXED-H 8*µ*m columns and a refractive index detector (Shodex RI-101) was used to determine molar mass distributions. Calibration with a series of near-monodisperse poly(ethylene oxide) (PEO) standards (with M*n* values ranging from 1.4 to 1,150 kDa). Samples were prepared at a concentration of 5 mg/mL in the phosphate buffer eluent and filtered (0.22 */mu*m, nylon). Dynamic light scattering (DLS) was performed using a Zetasizer Ultra (Malvern instruments, UK) with a cumulative fit correlation at 25^◦^C. The hydrodynamic diameter (D*_h_*), polydispersity index (PDI) and the *ζ*-potential were calculated using the ZS Xplorer software V3.1.064 (Malvern Panalytical Ltd., UK). The polymer was diluted to 0.25% w/w in filtered (0.22*µ*m nylon syringe filters) PBS and loaded in a Malvern Panalytical Folded Capillary Zeta Cells (DTS1070, Malvern). The acquired data were averaged over three consecutive runs and the D*_h_* values were reported from the obtained Z-Average.

#### Synthesis of 4-arm ZPs

4-arm CTA (synthesised in-house)[45], PSS, PVB and 4, 4’-azobis (4-cyanovaleric acid) (ACVA) (SI Table 1) were added to 0.1M sodium hydroxide solution (6 mL) and stirred at 65^◦^C under a nitrogen atmosphere for 20 hrs. Reaction mixture was oxygen-quenched under iced conditions. The product was diluted in deionised water and purified by dialysis (1000 MWCO) (Spectrum Labs) against Milli-Q water for 3 days (3 water changes/day). Samples were freeze-dried to produce a pale yellow powder.

**ZP12**: ^1^H NMR [400Mhz, (*D*_2_O)] *δ*=1.43 (s, 6*n* H); 2.88 (s, 9*n* H); 4.21 (s, 2*n* H); 6.61 (s, 4*n* H); 7.51 (s, 4*n* H) ppm. GPC: M*n* = 27 kDa, PDI = 2.19. DLS: *ζ*-potential = -29.1 mV, D*_h_* = 34.5 nm, PDI = 0.3. **ZP47**: ^1^H NMR [400Mhz, (*D*_2_O)] *δ*=1.54(s, 6*n* H); 2.91(s, 9*n* H); 4.28(s, 2*n* H); 6.70(s, 4*n* H); 7.52(s, 4*n* H) ppm. GPC: M*n* = 25 kDa, PDI = 2.09. DLS: *ζ*-potential = -29.1 mV, D*_h_* = 28.6 nm, PDI = 0.3. **ZP97**: ^1^H NMR [400Mhz, (*D*_2_O)] *δ* = 1.57 (s, 6*n* H); 2.97 (s, 9*n* H); 4.39 (s, 2*n* H); 6.66 (s, 4*n* H); 7.22 (s, 4*n* H) ppm; with *n* representing repeating units of the polymer. DLS: *ζ*-potential = 21.5 mV, D*_h_* = 21.5 nm, PDI = 0.6.

## Acknowledgements

The authors would like to thank Dr Siew-Wai Fong, Dr Hsiu-Yi Chen, Dr Nathan Wong and Dr Christopher Blanford for critical discussions and valuable suggestions on the study. We thank Elana Super and Dr Jungyeon Kim for providing the CTA core, and polymer supplies, respectively. We thank the members of the Jones Lab, Microbial Immunity Lab and Pathogen Modulation Lab for their continuous support throughout this project. We also thank A^∗^IDL Flow Cytometry platform and A^∗^IDL Mouse Core for assistance with cytometry analyses and support in animal breeding, respectively. Fig.1 was created using BioRender.com software.

## List of supplementary materials

Materials and Methods, SI Table 1 and SI Fig.1-14.

## Author contributions

S-L.M Data Curation, Design, Formal Analysis, Investigation, Visualisation, All Methodology, Writing – Original Draft, Writing – Review Editing. A.X.Y.L *In Vivo* Data Curation,Formal Analysis, Writing – Review Editing. M.A.C Maria A. Castillo: GPC and DLS Characterisation, Writing – Review Editing. L.J.B Data Curation for *in vitro* HSV-2, Writing – Review Editing. F-M.L *In Vivo* Conceptualisation, Formal Analysis, Visualisation, *In Vivo* Methodology, Writing – Review Editing. A.T-R *In Vivo* Conceptualisation, Formal Analysis, Visualisation, *In Vivo* Methodology. L.F.P.N *In vivo* Conceptualisation, Design, Funding Acquisition, Supervision Writing – Review Editing. S.T.J Polymer Conceptualisation, Design, Visualisation, Supervision, Writing – Original Draft, Writing – Review Editing. All authors revised and approved the final version of the manuscript.

## Data and materials availability

All data are available in the manuscript or the supplementary materials.

