## Supplementary Information for "A broad-spectrum, biocompatible, virucidal polymer reduces chikungunya virus in murine models"

### Supporting Information Available

#### Methods and Materials

Circulating leukocyte profiling

Briefly, 10 $\mu$L of blood from each mouse was collected from the tail and mixed in 10 $\mu$L of PBS supplemented with 10 $\mu$L citrate. The blood underwent lysis using the FC lysis buffer solution (R&D Systems). Live cells were stained with a Live/Dead Fixable Aqua dye before staining with the following cell-specific markers following manufacturer's protocol: BUV395-conjugated anti-CD45 (clone 30- F11), BV786-conjugated anti-CD3 (clone 17A2), BV421-conjugated anti-CD8 (clone 53-6.7), BV421-conjugated anti-B220 (clone 53-6.7), Phycoerythrin (PE)-conjugated anti-CD4 (clone GK1.5), BV650-conjugated anti-CD11b (clone M1/70), PE-CF594-conjugated anti-Ly6G (clone 1A8), APC/Fire 750-conjugated anti-Ly6C (HK1.4), APC-conjugated anti-NK1.1 (clone PK136) and Alexa Flour 700 (AF700)-conjugated anti-MHC-II (clone M5/114.15.2) for 30 min. All antibodies used for staining were purchased from BD Biosciences, BioLegend or ThermoFisher Scientific. Stained cells were fixed with IC Fixation Buffer (Invitrogen) and acquired using the NovoCyte Penteon (NovoExpress software, v4.0.5) and analysed with FlowJo v10.6.2 (Becton, Dickinson and Company).

Intracellular staining (ICS) of isolated $CD4^+$ T-cells

Spleens of naïve mice were harvested, homogenised and RBCs were lysed (as described above). The spleenocytes were subjected to $CD4^+$ T cell negative isolation using a MACS column loaded with anti-mouse $CD4^+$ microbeads, according to the manufacturer's instructions (Miltenyi Biotec). Efficiency of depletion was  85% as verified by flow cytometry. Following isolation, $CD4^+$ T cells were incubated overnight with various ZP12 concentrations (0-195 $\mu$g/mL). Hereafter, cells were incubated in complete IMDM (10% FBS, 1% PS) with or without PMA (40 ng/mL, Sigma-Aldrich) and 1 $\mu$g/mL Lonomycin (Sigma-Aldrich) at 37°C. After 1 hr, 2x GolgiPlug (Brefeldin A, Bioscience) and GolgiStop (Monensin, Bioscience), was added to the cells and incubated for a further 3 hrs[**?** ]. Cells were collected, washed, and stained with Live/Dead Fixable Aqua dye (Invitrogen) for 10 min, followed by surface marker staining for 20 min: PB-conjugated anti-CD4 (clone GK1.5, 1:400), BUV395-conjugated anti-CD25 (clone PC61, 1:200), APC-Cy7-conjugated anti-CD44 (clone IM7, 1:200), FITC-conjugated anti-CD69 (clone H1.2F3, 1:200) and PeCy7-conjugated anti-ICOS (clone C398.4A, 1:200). Cells were then fixed in fixation/permabilisation solution (FOXP3 kit, Invitrogen) as per the manufacturers instructions. Subsequently intracellular staining with IFN-$\gamma$ (APC, 1:200), IL-4 (PE, 1:200), TNF-$\alpha$ (BV650, 1:200) was performed. After washing cells, acquisition was done using the BD FACSymphony A5 (BD FACS Diva software v8.0; BD BioScience) and analysed with FlowJo v10.6.2 (Becton, Dickinson and Company).

Cytokine quantification by multiplexed bead-based immunoassays

Spleenocytes were incubated overnight with ZP12 in complete IMDM with or without PMA/lonomycin (T-cell stimulation), and with or without LPS (macrophage stimulation) (100 ng/mL, Sigma-Aldrich). The supernatants were collected, and immune mediators quantified using the LEGENDplex Mouse Inflammation Panel (13-plex) immunoassay (Biolegend), as per the manusfacturers protocol. The chemokines and cytokines assays included IL-23, IL-1α, IFN-γ, TNF-α, CCL2 (MCP-1), IL-12p70, IL-1β, IL-10, IL-6, IL-27, IL-17A, IFN-β and GM-CSF. Data were acquired using the NovoCyte Penteon (NovoExpress software, v4.0.5) and analysed using BioPlex Manager 6.0 software (Bio-Rad) based on standard curves plotted through a five-parameter logistic curve setting. Quantities of detected immune mediators are presented with a heat map using $\log_{10}$ concentration values scaled between 0 and 3 for visualisation.

Protein Corona Studies using PSS and ZP12

Four-week old male C57BL/6J mice were used for the *in vivo* intravenous (IV) PSS and ZP12 protein corona studies. CHIKV infection in mice were performed by inoculating $10^6$ PFUs CHIKV (in 30 $\mu$L PBS) subcutaneously in the ventral side of the right hind footpad near the ankle. Following 1 hr post-infection, 100 $\mu$L of 1 mg/kg ZP12 and 1 mg/kg PSS (other doses: 5 mg/kg ZP12) was administered via IV every 24 hrs for 7 days. For the non-treated groups, 100 $\mu$L PBS was administered. Joint swelling was measured as previously described.

SI Table 1: Values for different polymers and reagents. Key: CTA-Chain Transfer Agent. ACVA-4, 4'- azobis (4-cyanovaleric acid).

| **Polymer** | **ZP12** | | **ZP47** | | **ZP97** | |
|---|---|---|---|---|---|---|
| **Reagent** | **m (g)** | **n (mmol)** | **m (g)** | **n (mmol)** | **m (g)** | **n (mmol)** |
| CTA | 0.0262 | 0.0243 | 0.0262 | 0.0243 | 0.0262 | 0.0243 |
| ACVA | 0.0068 | 0.00242 | 0.0068 | 0.00242 | 0.0068 | 0.00242 |
| PSS | 0.90 | 4.37 | 0.70 | 3.39 | 0.30 | 1.45 |
| PVB | 0.10 | 0.49 | 0.31 | 1.45 | 0.72 | 3.39 |

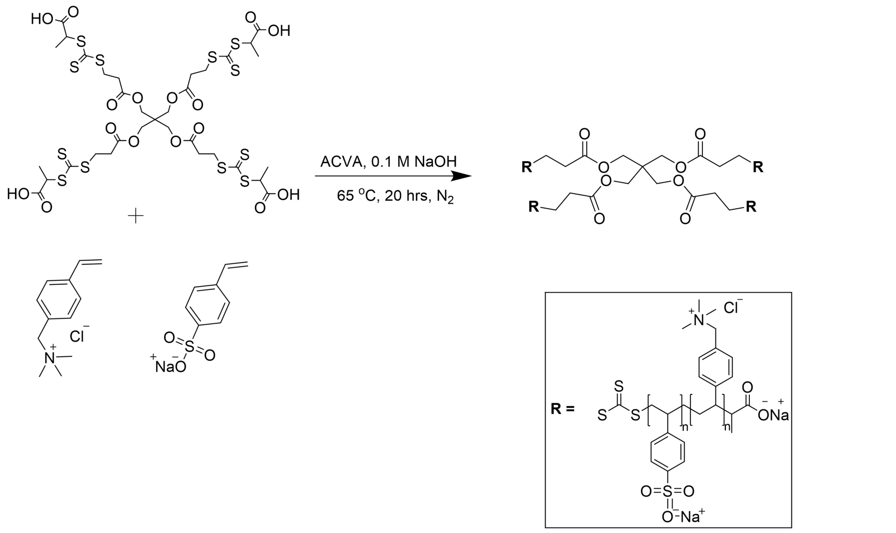


SI Fig.1: Chemical structure of CTA core (top) (with R depicting the polymeric arms attached to the core), monomers PVB (left) and PSS (right) for the synthesis of 4-arm star polymer.

**(a)**

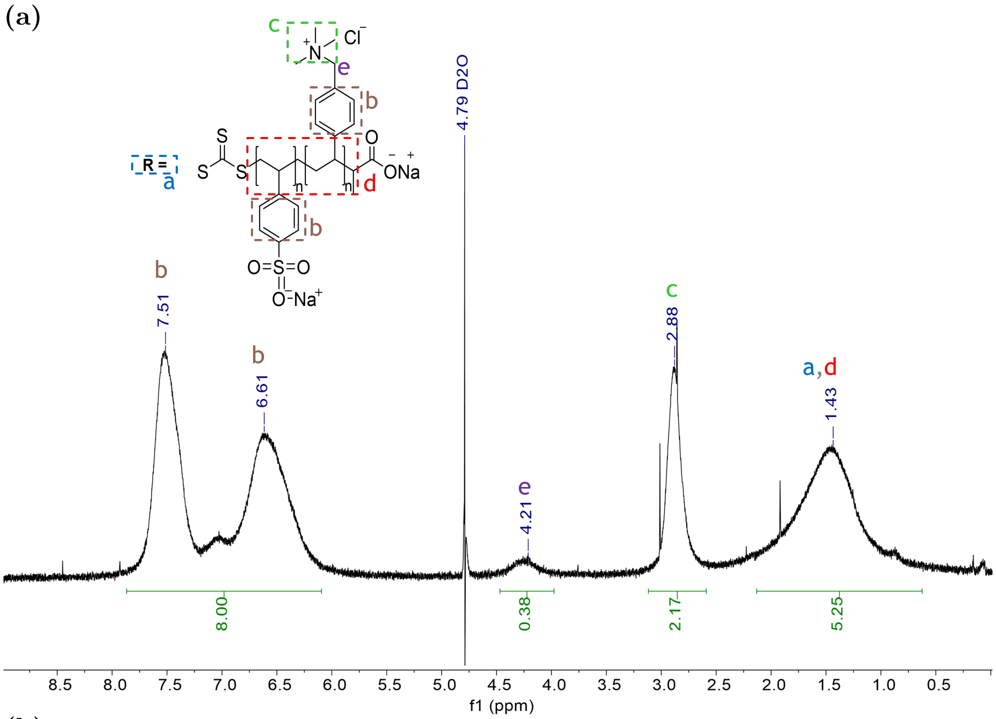

R =
a
S
S
S
n
n
d
O
ONa
b
b
c
N
Cl
e
O=S=O
O Na
4.79 D2O
b
7.51
b
6.61
e
4.21
c
2.88
a,d
1.43
8.00
0.38
2.17
5.25
8.5 8.0 7.5 7.0 6.5 6.0 5.5 5.0 4.5 4.0 3.5 3.0 2.5 2.0 1.5 1.0 0.5
f1 (ppm)


**(b)**

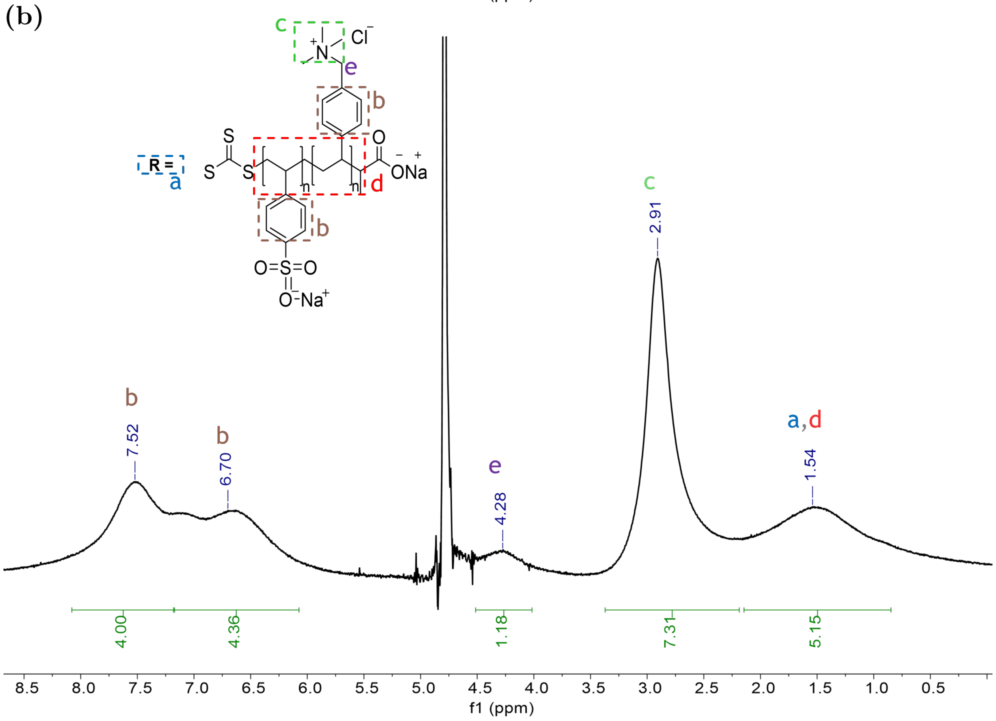

R =
a
S
S
S
n
n
d
O
ONa
b
b
c
N
Cl
e
O=S=O
O Na
b
7.52
b
6.70
e
4.28
c
2.91
a,d
1.54
4.00
4.36
1.18
7.31
5.15
8.5 8.0 7.5 7.0 6.5 6.0 5.5 5.0 4.5 4.0 3.5 3.0 2.5 2.0 1.5 1.0 0.5
f1 (ppm)

**(c)**

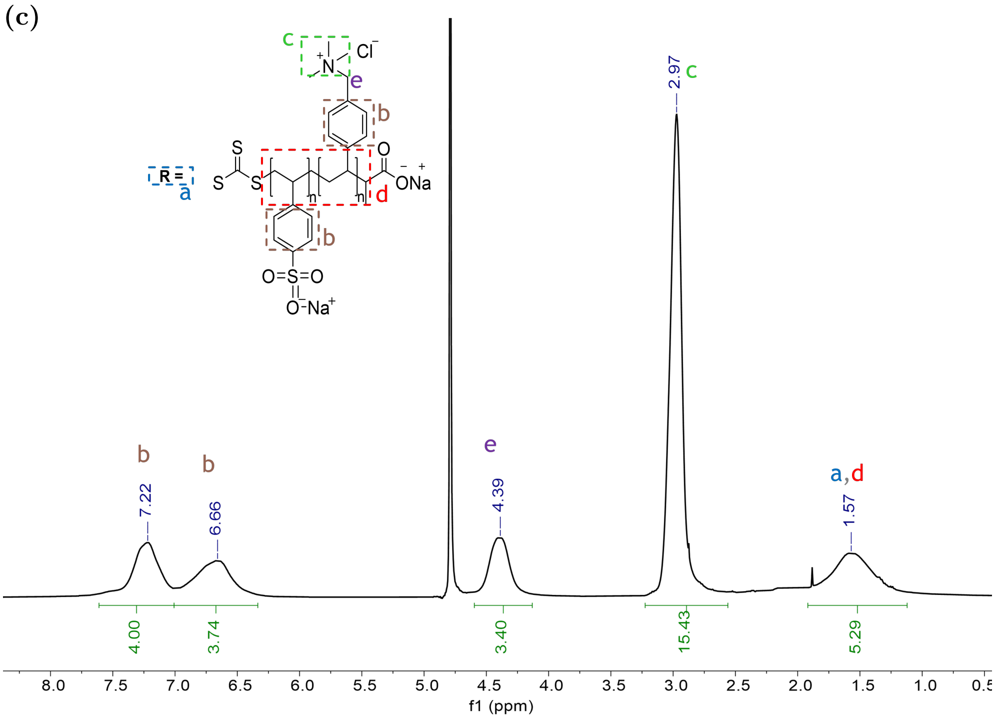


SI Fig.2: $^1$H NMR spectra of 4-arm DP50 Zwitterionic Polymers (ZPs) in $d_2O$ **(a)** ZP12 (12% PVB and 88% PSS) with the chemical structure showing proton environments (R depicting the growing monomer). Mw = 42.5 kDa. **(b)** ZP47 (47% PVB and 53% PSS). Mw = 42.8 kDa. **(c)** ZP97 (97% PVB and 3% PSS). Mw = 43.4 kDa.

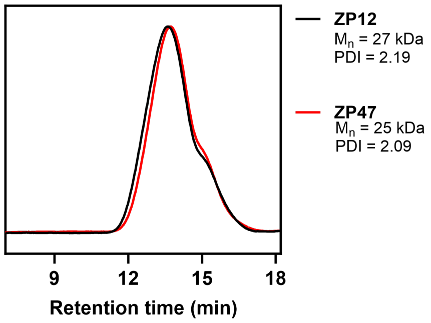


SI Fig.3: GPC trace for ZPs. Retention time for ZP12 ($M_n$: 27 kDa, PDI: 2.19) and ZP47 ($M_n$: 25 kDa, PDI: 2.09). Key: PDI: polydispersity ($M_w/M_n$).

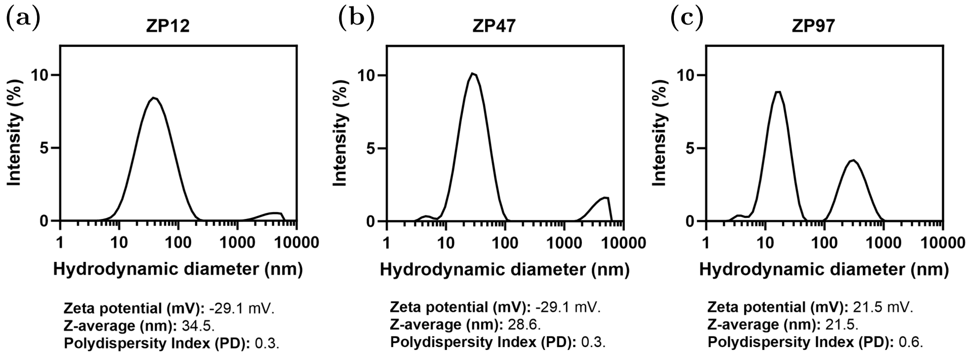


SI Fig.4: Dynamic Light Scattering (DLS) spectra for ZPs. **(a)** ZP12 ($\zeta$-potential = -29.1 mV, $D_h$= 34.5 nm, PDI = 0.3). **(b)** ZP47 ($\zeta$-potential = -29.1 mV, $D_h$= 28.6 nm, PDI = 0.3). **(c)** ZP97 ($\zeta$-potential = 21.5 mV, $D_h$= 21.5 nm, PDI = 0.6).

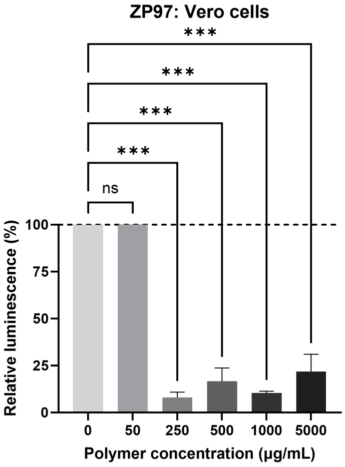


SI Fig.5: Vero-E6 cell viability of ZP97 assessed using the Cell Titre Glo assay. Percent of viable cells were calculated based on untreated (PBS) cells. Data are presented as means±SEM of three independent experiments performed in triplicate. Statistical analyses were performed using one-way ANOVA (***p<0.001).

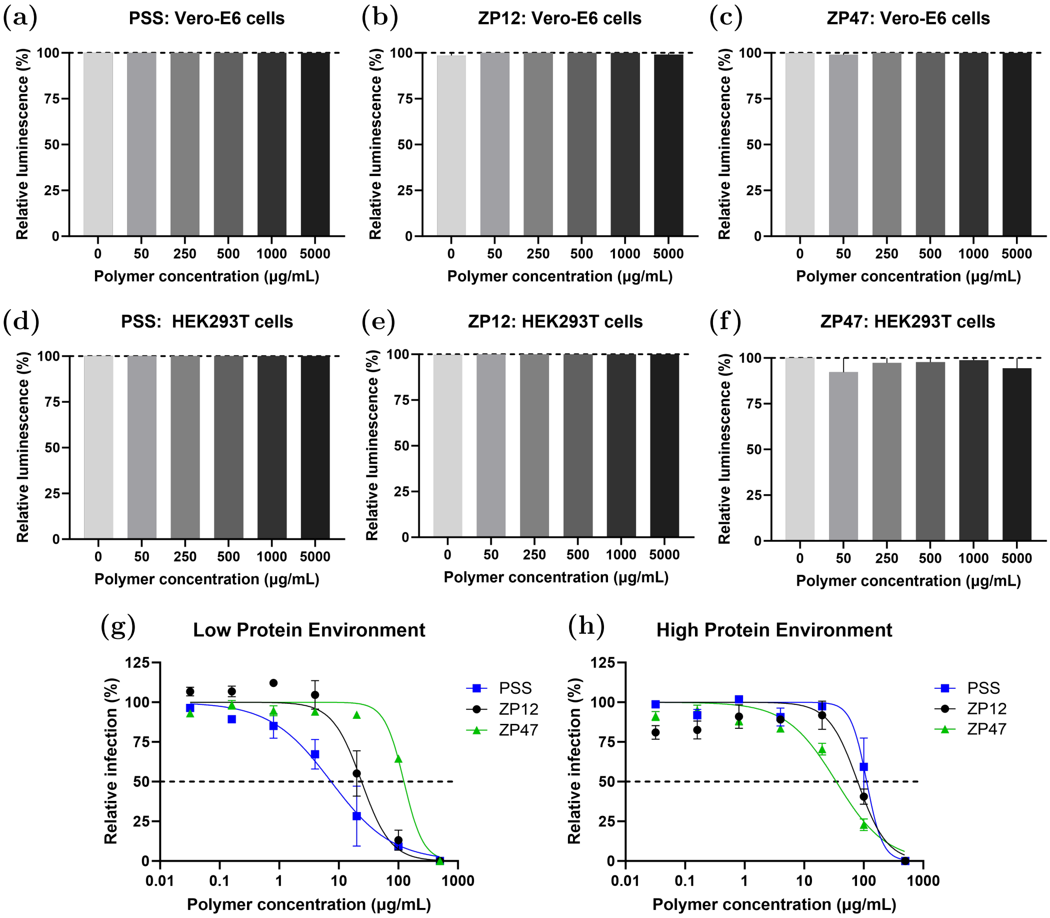


SI Fig.6: Toxicity and antiviral activities of PSS, ZP12 and ZP47 against CHIKV. Cell viability assessed using the Cell Titre Glo assay at various polymer concentrations ranging from 50 $\mu$g/mL to 5000 $\mu$g/mL. **(a-c)** Vero-E6 cells and **(d-f)** HEK293T cells. Percent of viable cells were calculated based on untreated (PBS) cells. Dose response curves of PSS (blue square trace), ZP12 (black dot trace) and ZP47 (green triangle trace) in **(g)** a low protein environment and **(h)** a high protein environment (0.01% BSA) showing $IC_{50}$ (dotted line). Percentage infection were expressed relative to signals from untreated infected cells. Data are presented as means±SEM of three independent experiments performed in at least duplicate.

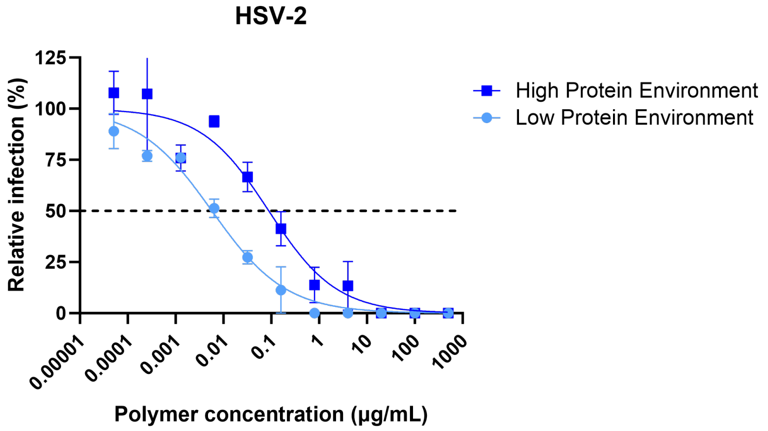


SI Fig.7: Antiviral activity of star-PSS against HSV-2 (MOI 1) in a low and high protein environment. Dose response curve showing PSS $IC_{50}$ (dotted line). High protein environment: $IC_{50}$= 0.0021 $\mu$M and Low protein environment: $IC_{50}$=0.000136 $\mu$M. Percentage infection were expressed relative to plaques from untreated infected cells. Data are presented as means±SEM of three independent experiments performed in duplicate.

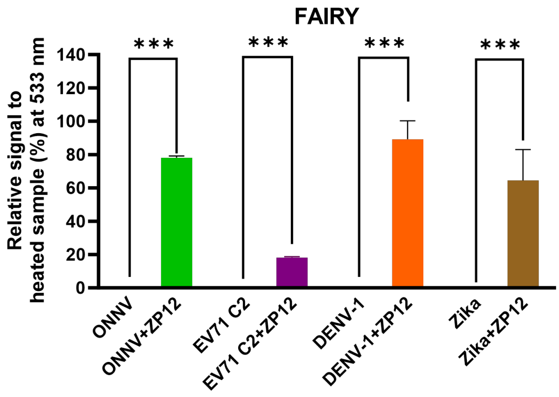


SI Fig.8: FAIRY assay demonstrating broad-spectrum virucidal properties of ZP12 against a range of viral families. Signals are normalised to 10 min 90°C heated samples. Data are presented as means±SEM of three independent experiments performed in triplicate.

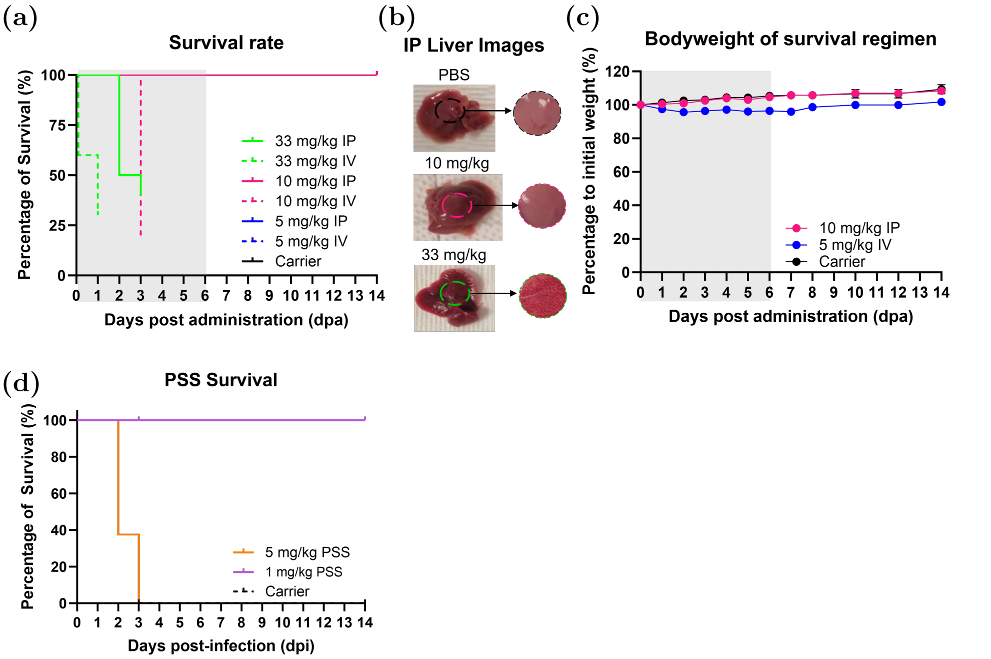


SI Fig.9: Biocompatability of ZP12 (intraperitoneal - IP and intravenous - IV) and PSS (IV). **(a)** Survival curve comparing different ZP12 regimen doses between IP and IV. We found that 33 mg/kg was toxic to the animals with both routes. For the IP route, 10 mg/kg was well tolerated, whilst 5 mg/kg was well-tolerated for the IV route. **(b)** Images of liver following intraperitoneal administration with zoomed regions. **(c)** Bodyweight of animals with highest tolerable dose for IP and IV. **(d)** Survival curve comparing different PSS doses when administered IV. We found that 5 mg/kg was toxic to the animals. Survival curves analysed using Mantel-cox test. Data are presented as means±SEM of at least 5 animals per experimental group and are representative of two independent experiments.

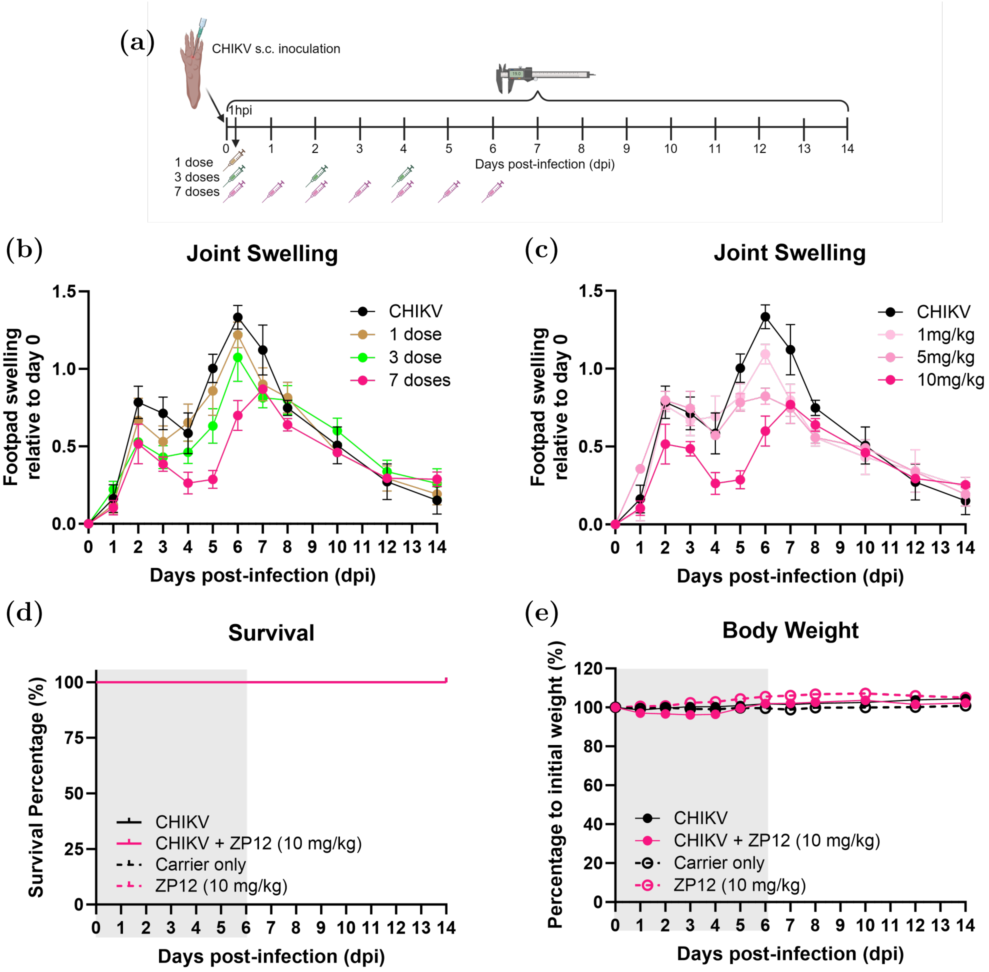


SI Fig.10: ZP12 regimen investigation against CHIKV-infected C57BL/6 murine models. **(a)** Schematic representing virus challenge and 1 hours post infection (hpi) treatment regimen of ZP12 administered. **(b)** Joint swelling of animals that were treated with 10 mg/kg ZP12 at different frequencies: 1 dose, 3 doses (dose every 48 hours) and 7 doses (dose every 24 hours for 7 days). **(c)** Joint swelling of animals that received 7 doses of ZP12 at different concentrations: 1 mg/kg, 5 mg/kg and 10 mg/kg. **(d-e)** Survival curve analysed using Mantel-cox test and body weight of CHIKV-infected animals treated with 10 mg/kg ZP12 every 24 hours for 7 days. All data are presented as mean±SEM of at least 3 animals per experimental group and are representative of two independent experiments.

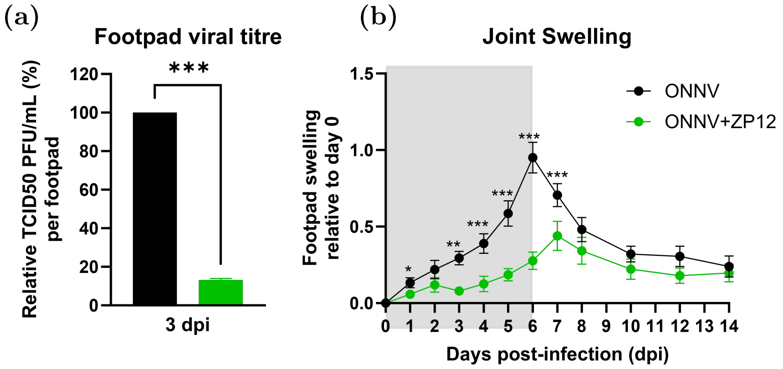


SI Fig.11: ONNV-infected C57BL/6 murine models treated with 10 mg/kg ZP12 via IP every 24 hrs for 7 days. **(a)** Footpad tissue viral load. **(b)** Joint swelling. All data are presented as mean±SEM of at least 5 animals per experimental group and are representative of two independent experiments.

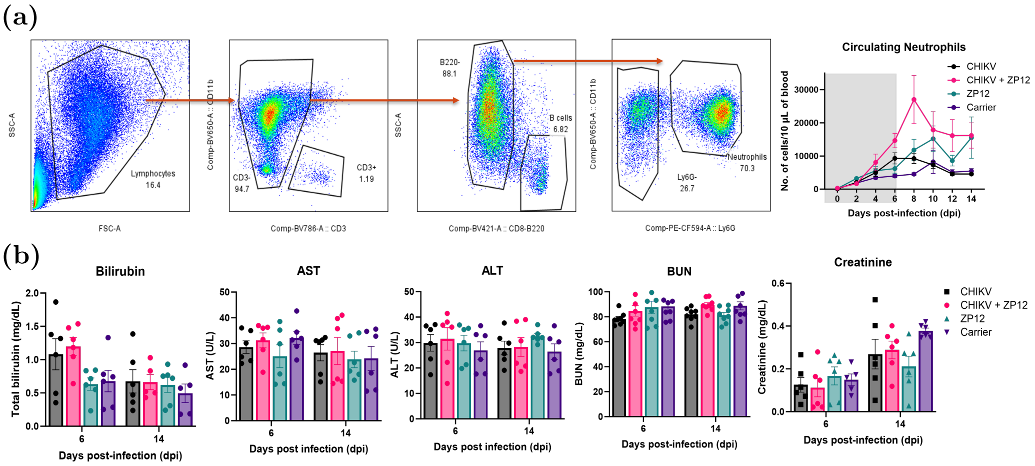


SI Fig.12: Investigating increased circulating neutrophils. **(a)** Gating strategy. **(b)** Liver (bilirubin, AST, ALT) and kidney (BUN, creatinine) metabolism assessment. Data are presented as mean±SEM of at least 3 animals per experimental group from two independent experiments.

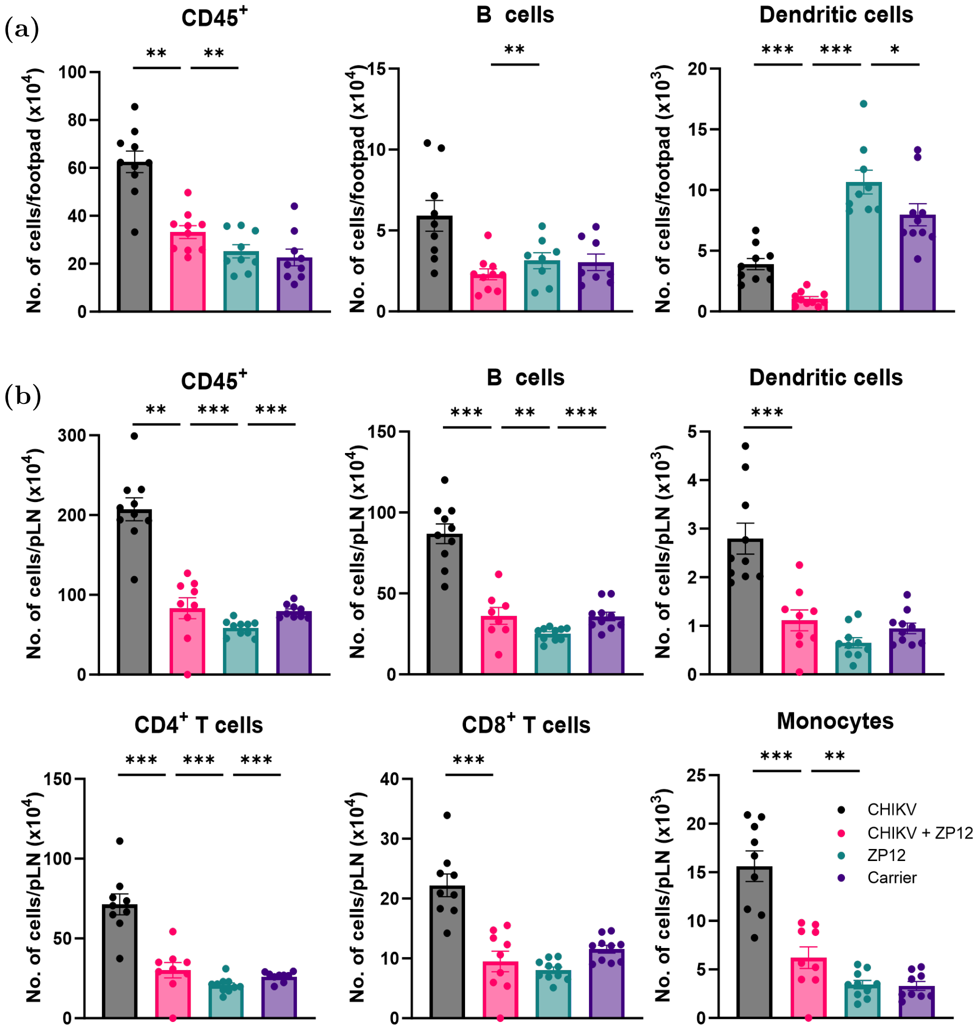


SI Fig.13: Leukocytes in the **(a)** footpad and **(b)** popliteal lymph node (pLN) reveal decreased joint immune profile of infiltrating immune cells of CHIKV-ZP12 treated animals. Data are presented as mean±SEM of at least 5 animals per experimental group and are representative of two independent experiments.

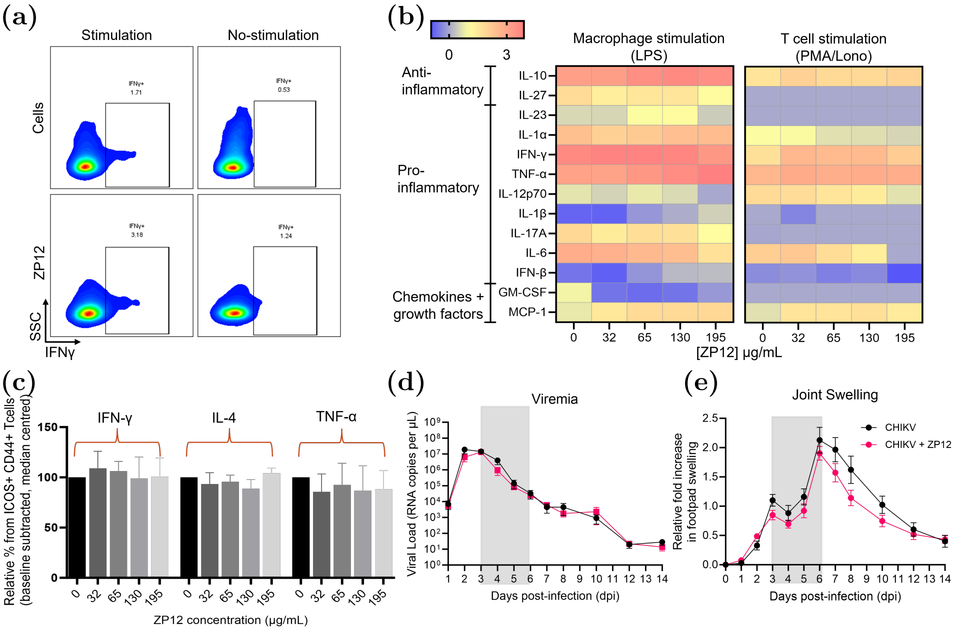


SI Fig.14: Investigating if ZP12 is an immuno-supressant. **(a,c)**The isolated $CD4^+$ T cells were incubated overnight with ZP12 and then stimulated for 4 h with PMA and ionomycin before intracellular antibody staining. **(a)** Representative flow cytometry plot for SSC and IFNγ staining in ZP12 with and without stimulation. **(c)** Median-centred proportion of $ICOS^+CD44^+CD4^+$ T cells across two independent experiments ($n = 5$ per group). **(b)** Spleenocyte cells were incubated overnight with ZP12 and then either with PMA and ionomycin (T-cells) or lipopolysaccharide (LPS) (macrophage) before a multiplex microbead-based assay was used to quantify the levels of immune mediators present in these samples. Heat map showing the levels of the analysed immune mediators in each group of animals. **(d-e)** CHIKV infected animals were treated with ZP12 3 dpi every 24 hours for 3 days. **(d)** viremia and **(e)** joint swelling. Data presented were obtained from at least two independent experiments. All data are presented as mean±SEM. Comparison between ZP12 and PBS groups were performed with non-parametric Mann-Whitney U test.
